# SuperCRUNCH: A bioinformatics toolkit for creating and manipulating supermatrices and other large phylogenetic datasets

**DOI:** 10.1101/538728

**Authors:** Daniel M. Portik, John J. Wiens

## Abstract

1. Phylogenies with extensive taxon sampling have become indispensable for many types of ecological and evolutionary studies. Many large-scale trees are based on a “supermatrix” approach, which involves amalgamating thousands of published sequences for a group. Constructing up-to-date supermatrices can be challenging, especially as new sequences may become available almost constantly. Additionally, genomic datasets (composed of thousands of loci) are becoming common in phylogenetics and phylogeography, and present novel challenges for constructing such datasets.
2. Here we present SuperCRUNCH, a Python toolkit for assembling large phylogenetic datasets. It can be applied to GenBank sequences, unpublished sequences, or combinations of GenBank and unpublished data. SuperCRUNCH constructs local databases and uses them to conduct rapid searches for user-specified sets of taxa and loci. Sequences are parsed into putative loci and passed through rigorous filtering steps. A post-filtering step allows for selection of one sequence per taxon (i.e. species-level supermatrix) or retention of all sequences per taxon (i.e. population-level dataset). Importantly, SuperCRUNCH can generate “vouchered” population-level datasets, in which voucher information is used to generate multi-locus phylogeographic datasets. SuperCRUNCH offers many options for taxonomy resolution, similarity filtering, sequence selection, alignment, and file manipulation.
3. We demonstrate the range of features available in SuperCRUNCH by generating a variety of phylogenetic datasets. Output datasets include traditional species-level supermatrices, large-scale phylogenomic matrices, and phylogeographic datasets. Finally, we briefly compare the ability of SuperCRUNCH to construct species-level supermatrices to alternative approaches. SuperCRUNCH generated a large-scale supermatrix (1,400 taxa and 66 loci) from 16GB of GenBank data in ∼1.5 hours, and generated population-level datasets (<350 samples, <10 loci) in <1 minute. It outperformed alternative methods for supermatrix construction in terms of taxa, loci, and sequences recovered.
4. SuperCRUNCH is a modular bioinformatics toolkit that can be used to assemble datasets for any taxonomic group and scale (kingdoms to individuals). It allows rapid construction of supermatrices, greatly simplifying the process of updating large phylogenies with new data. It is also designed to produce population-level datasets. SuperCRUNCH streamlines the major tasks required to process phylogenetic data, including filtering, alignment, trimming, and formatting. SuperCRUNCH is open-source, documented, and available at https://github.com/dportik/SuperCRUNCH.

## 1 INTRODUCTION

Large-scale phylogenies, including hundreds or thousands of species, have become essential for many studies in ecology and evolutionary biology. Many of these large-scale phylogenies are based on the supermatrix approach (e.g., de Queiroz & Gatesy, 2007), which typically involves amalgamating thousands of sequences from public databases (e.g., GenBank). Several tools exist for assembling these datasets, which differ in their approach and functionality. These include programs like PhyLoTA (Sanderson, Boss, Chen, Cranston, & Wehe, 2008), PHLAWD (Smith, Beaulieu, & Donoghue, 2009), phyloGenerator (Pearse & Purvis, 2013), SUMAC (Freyman, 2015), SUPERSMART (Antonelli et al., 2017), PhylotaR (Bennett et al., 2018) and PyPHLAWD (Smith & Walker, 2018). These programs have greatly improved access to GenBank data, and several have been successfully used in a variety of research contexts. However, each program comes with its own pros and cons for assembling molecular datasets. For example, several programs employ automated (“all-by-all”) clustering of all sequences. This approach enables the discovery of orthologous sequence sets, which is useful if target loci are unknown. However, searching for specific loci is generally incompatible with the “all-by-all” clustering approaches. As genomic datasets composed of several thousand loci become increasingly common, the ability to specify target loci will be essential for building customized phylogenomic datasets. Additionally, many programs use a GenBank database release to obtain starting sequences, in which users specify a taxon and the relevant sequence data are downloaded automatically. This design is useful and convenient, but it unfortunately prevents the inclusion of locally generated (e.g., unpublished) sequence data, thereby limiting analyses to published sequences. A majority of methods were designed to create species-level datasets, in which a species is represented by one sequence per locus (e.g., a traditional supermatrix). It is typically not possible to use these methods to generate phylogeographic (population-level) datasets, in which a species is represented by many individuals sequenced for anywhere from one gene to thousands of loci. The dramatic increase in the availability and size of phylogeographic datasets (McCormack et al., 2013; Garrick et al., 2015) has created a need for methods which can construct large-scale population-level datasets. Additionally, no current methods utilize voucher codes (e.g., a field series, museum number, or other identifier). These codes are critical for linking samples and building phylogeographic datasets. Thus, producing high-quality phylogenetic datasets can be challenging using the available methods, and programs with new functionality are required to keep pace with the changing landscape of phylogenetics and phylogeography.

To address these challenges, we developed SuperCRUNCH, a semi-automated method for creating phylogenetic and phylogeographic datasets. SuperCRUNCH can be used to process sequences from GenBank, datasets containing only locally generated (unpublished) sequences, or a combination of sequence types. During initial steps, the sequence data are parsed into loci based on user-supplied lists of taxa and loci, offering fine-control for targeted searches. SuperCRUNCH allows any taxonomy to be used, and offers simple steps for identifying and resolving taxonomic conflicts. SuperCRUNCH also includes refined methods for similarity filtering, quality filtering, and sequence selection. By offering the option to select one representative sequence per species or retain all filtered sequences, SuperCRUNCH can be used to generate species-level datasets (one sequence per species per gene) and population-level datasets (multiple sequences per species per gene). SuperCRUNCH can also filter sequences using voucher codes, which can label and link sequences in phylogeographic datasets (e.g., a “vouchered” dataset). Analyses are highly scalable and can range in size from small population-level datasets (one taxon, one gene) to large phylogenomic datasets (hundreds of taxa, thousands of loci). SuperCRUNCH is modular in design, offering flexibility across all major steps in constructing phylogenetic datasets, and analyses are transparent and highly reproducible. SuperCRUNCH is open-source, heavily documented, and freely available at https://github.com/dportik/SuperCRUNCH.

## 2 INSTALLATION

SuperCRUNCH consists of a set of python modules that function as stand-alone command-line scripts. As of SuperCRUNCH v1.2, these modules can be run using Python 2.7 or 3.7. All modules can be downloaded and executed independently without the need to install SuperCRUNCH as a python package or library, making them easy to use and edit. Nevertheless, there are eight dependencies that should be installed that enable the use of all features in SuperCRUNCH. These include the biopython package for python, and the following seven external dependencies: ncbi-blast+ (for blastn and makeblastdb; Altschul, Gish, Miller, Myers, & Lipman, 1990; Camacho et al., 2009), cd-hit-est (Li & Godzik, 2006), clustal-o (Sievers et al., 2011), mafft (Katoh, Misawa, Kuma, & Miyata, 2002; Katoh & Standley, 2013), muscle (Edgar, 2004), macse (Ranwez, Douzery, Cambon, Chantret, & Delsuc, 2018), and trimal (Capella-Gutiérrez, Silla-Martínez, & Gabaldón, 2009). Installation instructions for all dependencies is provided on the SuperCRUNCH github wiki (https://github.com/dportik/SuperCRUNCH/wiki).

## 3 WORKFLOW

A comprehensive user-guide, including overviews for all major steps and detailed instructions for all modules, is available on the SuperCRUNCH github wiki. Several complete analyses are posted on the Open Science Framework SuperCRUNCH project page, available at: https://osf.io/bpt94. Here, we briefly outline the major steps in a typical analysis, including some technical details for key steps. However, we strongly encourage users to read the complete documentation available online.

### 3.1 Overview

SuperCRUNCH is designed to work with fasta-formatted sequence data that have been previously downloaded (e.g., from GenBank) or are locally available (e.g., processed sequences from in-house projects). No connection to live databases (such as NCBI) is required. Three input files are needed to perform a typical analysis: a set of sequence records in fasta format, a list of taxonomic names, and a list of loci (or genes) and associated search terms. The contents of these input files are described in greater detail below. The general workflow involves assessing taxonomy, parsing loci, similarity filtering, sequence selection, sequence alignment, and various post-alignment tasks (Fig. 1). The taxonomy used is user-supplied (e.g., not explicitly linked to any online databases). Therefore, an important first step is to identify and resolve potential conflicts between the user-supplied taxon list and the taxon labels in the sequence records. Afterwards, searches are conducted to identify records that putatively belong to loci (based on the content of record labels). These records are then written to locus-specific files. The sequences in each locus are then subjected to more stringent filtering using similarity searching (via nucleotide BLAST). This step removes non-homologous sequences and trims homologous sequences to remove non-target regions. After similarity filtering, the sequence-selection step allows selection of one sequence per species per locus or including all sequences. For both options, several additional filters (e.g., requiring an error-free reading frame, minimum length, or voucher information) can be used to ensure only high-quality sequences are retained. Sequences can then be prepared for alignment (adjusting direction and/or reading frame) and subsequently aligned using several alignment methods. After alignment, sequences can be relabeled, and the alignments can be trimmed, converted to multiple formats, and concatenated. SuperCRUNCH analyses end with the production of fully formatted input files that are compatible with numerous phylogenetic and population-genetic programs. Below, we provide additional details for the major steps outlined here.

**FIGURE 1.**
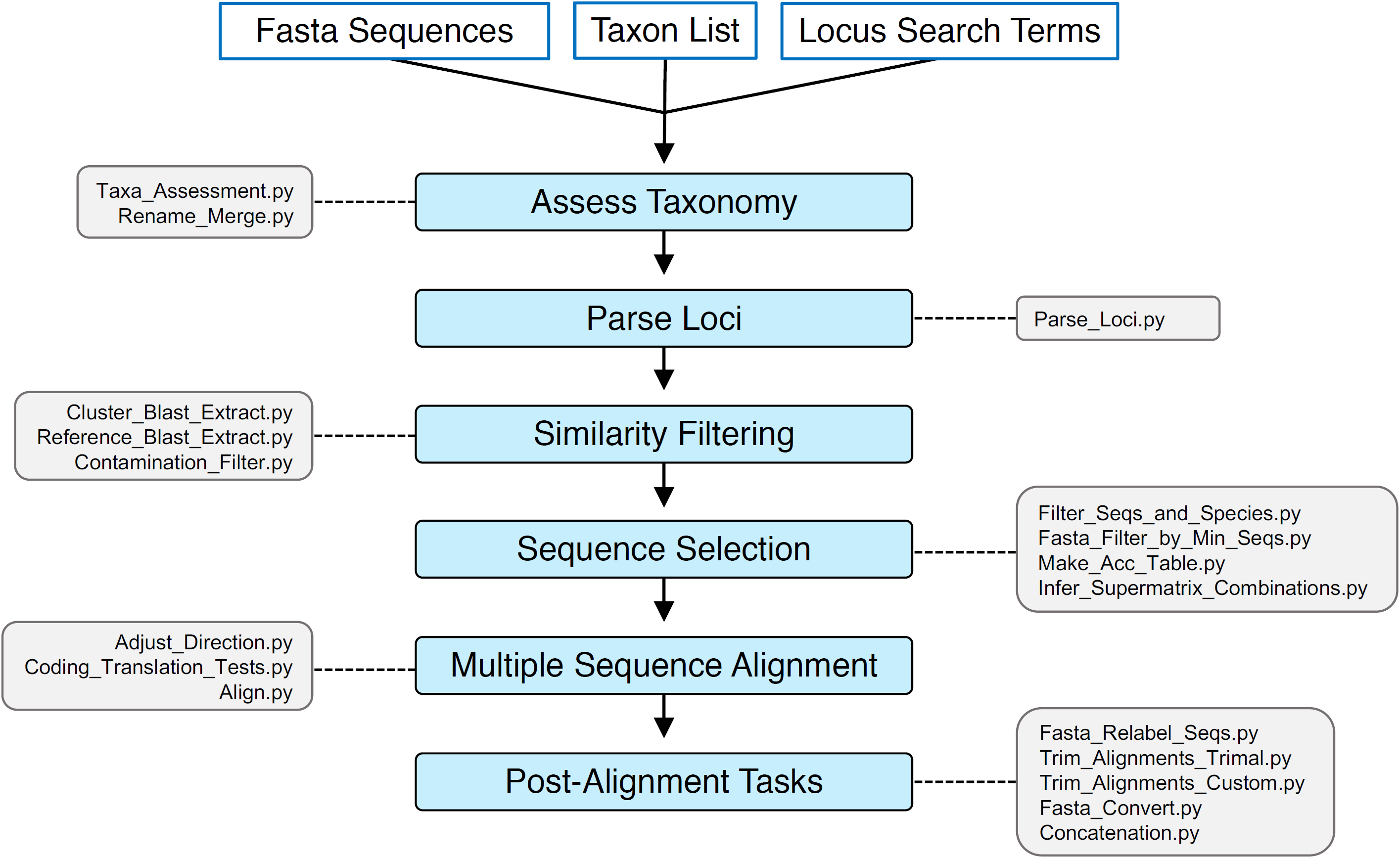
A depiction of the general steps (and associated modules) involved in full SuperCRUNCH analyses. Each step is outlined in a corresponding entry of the same title in the Workflow section of the main text.

### 3.2 Starting Sequences

SuperCRUNCH requires a single fasta file of nucleotide sequence records as the main input. The fasta file can contain records from GenBank, unpublished sequence records, or a combination. GenBank data can be obtained by searching for relevant taxonomy terms or organism identifier codes on that database, and downloading the records in fasta format. For clades with many species, downloading all records directly may not be possible. For these groups, results from multiple searches using key organism identifiers can be downloaded and combined into a single fasta file. Automated downloading of GenBank sequence data through SuperCRUNCH is currently not supported, but will be included in a future release. Locally generated data should be formatted similar to GenBank records. A typical record should contain an accession number (a unique identifier code), a taxon label (two-part or three-part name, genus/species or genus/species/subspecies), and locus information (gene abbreviation and/or full name). Voucher information is optional. Additional details and examples of how to label Sanger-sequenced and sequence-capture datasets are provided in the online documentation.

### 3.3 Assessing Taxonomy

SuperCRUNCH allows any taxonomy to be used. Taxonomy is supplied as a simple text file with one taxon name per line. Two-part and three-part names can be used. SuperCRUNCH offers the option to include or exclude subspecies. If subspecies are excluded, the third component of any three-part name is ignored, thereby reducing it to a two-part name. A taxon list can therefore contain a mix of species and subspecies names, even if subspecies are not desired. Although SuperCRUNCH does not connect with any taxonomy databases, lists of taxon names for large clades can be obtained through such databases, including the NCBI Taxonomy Browser or Global Names Database (Patterson et al., 2016). Many groups also have taxonomic databases, such as the Reptile Database (Uetz, Freed, & Hošek, 2018) and AmphibiaWeb (2019). These usually contain up-to-date taxonomies in a downloadable format. Taxon names can also be extracted directly from fasta files using the *Fasta_Get_Taxa.py* module. This option is most useful for unpublished sequences and sequence sets with few species.

Ideally, the user-supplied taxonomy will match the taxon names in the sequence records. However, taxonomy can change rapidly and conflicts often arise. To pass initial filtering steps, a record must have a taxon label that matches a name in the user-supplied taxonomy. Before beginning any filtering steps, it is therefore important to understand how compatible the user-supplied taxonomy is with the sequence-record set. The *Taxa_Assessment.py* module will perform an initial search across records to identify all records with a taxon label contained in the provided taxonomy, and identify all records with an unmatched taxon label (which would fail initial filtering steps). A list of unmatched taxon names is provided as output. External tools such as organismal databases, taxize/pytaxize (Chamberlain & Szöcs, 2013; Chamberlain et al., 2017), or the resolver function in the Global Names Database (Patterson et al., 2016), can be used to identify a “correct” name for an unmatched name. If a set of updated names is supplied for a set of unmatched names, the *Rename_Merge.py* module can be used to relabel all relevant records with the updated names, thus allowing them to pass the initial filtering steps. The combination of these two taxonomy modules allows users to correct minor labeling errors (such as misspellings), reconcile synonymies, or completely update names to a newer taxonomy.

### 3.4 Parsing Loci

The *Parse_Loci.py* module conducts searches for specific loci using a set of user-supplied search terms, including gene abbreviations and full gene names. All searches are conducted using SQL with a local database constructed from the input sequences, and the initial assignment of a sequence to a locus is based purely on matches to the record labeling. For a sequence to be written to a locus-specific file, it must match either the gene abbreviation or description for that locus, and it must have a taxon label present in the user-supplied taxonomy. This approach creates smaller locus-specific sequence sets from the initial sequence set, which are more tractable for downstream similarity searches (versus “all-by-all” clustering).

The success of finding sequences using SuperCRUNCH depends on providing appropriate gene abbreviations and labels. We recommend searching on GenBank to identify common labeling or using gene databases such as GeneCards (Stelzer et al., 2016). There is no hard upper bound on how many loci can be searched for. Thus, SuperCRUNCH can be used to process large phylogenomic datasets (e.g., sequence capture experiments) including those with thousands of species and loci. Whole mitochondrial genomes can also be included for any search involving a particular mitochondrial gene (see below). Recommendations for optimizing locus searches for different data types are provided in the online documentation.

The choice of loci will be group-specific. Previous phylogenetic/phylogeographic papers can be used to identify appropriate loci. The best criteria for selecting loci remain unresolved. One relevant criterion is completeness (e.g., including only loci present in >20% of the species). For each search conducted with *Parse_Loci.py*, the number of sequences found for each locus will be output. Therefore, it can be used to survey the availability of sequences for each locus. A downstream step allows loci to be filtered based on a minimum number of required sequences, so decisions can be made after additional filtering.

The *Parse_Loci.py* module performs another important task: automatically detecting voucher information in those sequence record labels containing a “voucher”, “strain”, or “isolate” field (see online documentation). This information is written into the records as a new tag that is discoverable in other downstream steps, allowing the creation of “vouchered” datasets.

### 3.5 Similarity Filtering

SuperCRUNCH offers two parallel methods for filtering sequences based on similarity. Each method uses nucleotide BLAST to perform searches, but they differ in whether reference sequences are automatically selected (*Cluster_Blast_Extract.py*) or user-provided (*Reference_Blast_Extract.py*) (Fig. 2). The automatic selection of reference sequences is appropriate for loci consisting of “simple” sequence records (Fig. 2). We define “simple” record sets as those generally containing a single gene region with limited length variation, which results from use of the same primers (Sanger-sequencing) or probes (sequence capture) to generate sequences. The *Cluster_Blast_Extract.py* module can be used for these types of loci. These generally include nuclear markers and those from commercial probe sets (e.g., UCEs: ultraconserved elements). The *Cluster_Blast_Extract.py* module begins by clustering sequences based on similarity using cd-hit-est. It then identifies the largest sequence cluster, and designates that as the reference sequence set (Fig. 2). All starting sequences (including those in the reference cluster) are then blasted to this reference using BLASTn. This method is convenient for automating the process of similarity filtering for “simple” records and can be used to screen thousands of loci.

**FIGURE 2.**
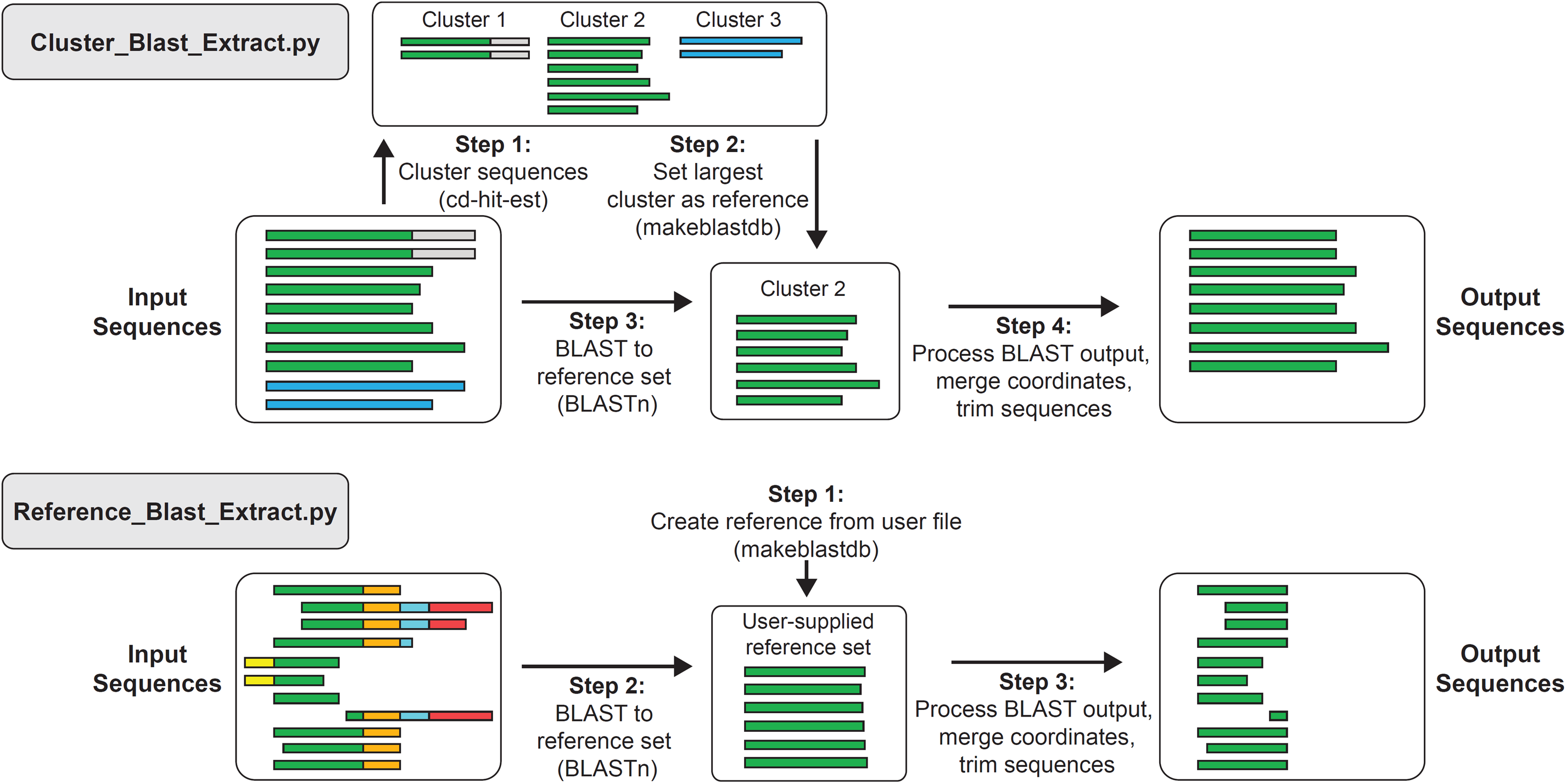
An illustration of the similarity searching workflows occurring in the *Cluster_Blast_Extract.py* and *Reference_Blast_Extract.py* modules. Green color represents target regions, and all other colors represent non-target regions.

However, *Cluster_Blast_Extract.py* will fail for loci containing “complex” sequence records. “Complex” records include those containing the target region plus non-target sequence (e.g., other regions or genes). Common examples include long mtDNA fragments and whole mitogenomes (Fig. 2). Another type of “complex” record is a gene sequenced for different fragments that have little or no overlap. For these sequence sets, the *Reference_Blast_Extract.py* module should be used instead. Rather than identifying the reference set from the starting sequences via clustering, it requires a user-supplied reference sequence set to perform BLASTn searches (Fig. 2). An external reference set must be provided for each locus, and it ensures that only the desired regions are targeted and extracted. For example, a set of ND2 reference sequences can be used to extract only ND2 regions from a record set comprised of whole mitochondrial genomes, multi-gene mitochondrial sequences, and partial ND2 records.

For both modules, the BLASTn algorithm can be specified by the user (blastn, blastn-short, megablast, or dc-megablast), allowing searches to be tailored to inter- or intraspecific datasets. After BLASTn searches are conducted for a locus, sequences without significant matches are discarded. For all other sequences, the BLAST coordinates of all hits (excluding self-hits) are merged to identify the target region of the query sequence. Based on these coordinates, the entire sequence or a trimmed portion of the sequence is kept. The BLAST coordinate merging action often results in a single continuous interval (e.g., bases 1–800). However, non-overlapping coordinates can also be produced (e.g., bases 1–200, 450–800). Two common examples (sequences containing stretches of N’s or gene duplications) are illustrated in Figure 3.

**FIGURE 3.**
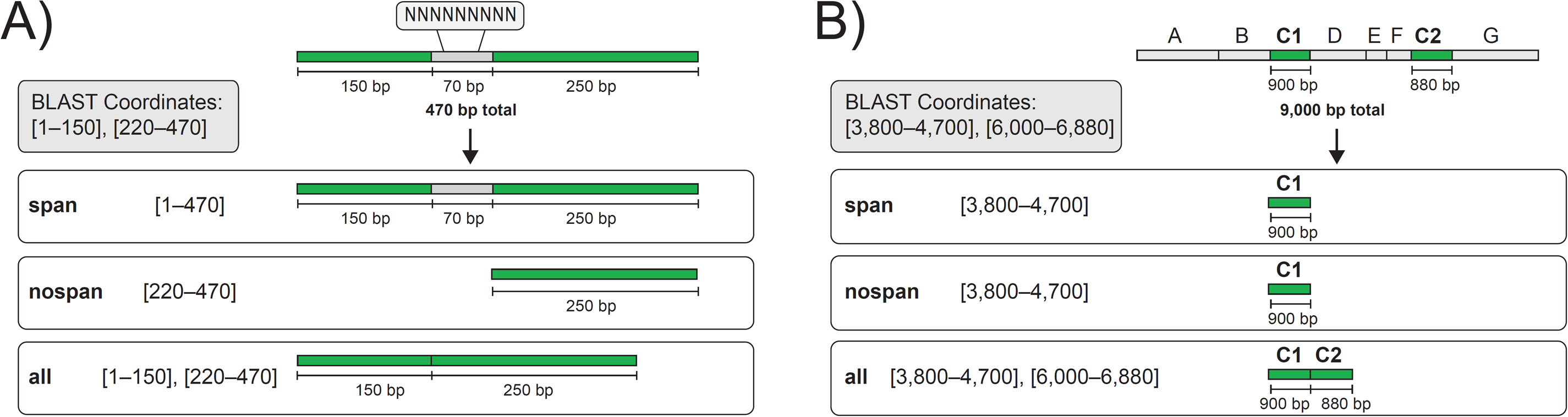
A demonstration of the options available for handling non-overlapping BLAST coordinates for query sequences with two common examples: (A) a sequence that contains a stretch of N’s, and (B) a long sequence containing multiple genes (represented by letters) that also contains a gene duplication (indicated by C1 and C2), such as an organellar genome. In both sequences, green represents the target region and grey represents either missing data (A) or non-target regions (B). The resulting merged BLAST coordinates are shown for each sequence, along with which coordinates would be selected under the available options (“span”, “nospan”, and “all”, see main text).

Multiple options are available for handling non-overlapping sequence intervals. The default option is “span”, which bridges non-overlapping intervals <X base pairs apart, where X is the default value (100 bp) or a user-supplied value. However, if the gap is >X bases, the longest interval is selected instead. The “nospan” method will simply select the longest interval of the coordinate set, and the “all” method will concatenate the sequence intervals together. Results from each option are shown in Figure 3.

An optional contamination-filtering step is available (*Contamination_Filter.py*). This step excludes all sequences scoring >95% identity for at least 100 continuous base pairs to the reference sequences. Here, the contamination reference sequences should correspond to the expected source of contamination (see documentation).

### 3.6 Sequence Selection

SuperCRUNCH can construct two fundamentally different datasets: species-level supermatrices and population-level (phylogeographic) datasets. The *Filter_Seqs_and_Species.py* module is used to select the sequences necessary to construct either dataset (using the “oneseq” or “allseqs” options). For supermatrices, a single sequence is typically used to represent each species for each gene. If multiple sequences for a given gene exist for a given species (e.g., because multiple individuals were sampled), then an objective strategy must be used for sequence selection. *Filter_Seqs_and_Species.py* offers several options, including the simplest solution: sorting sequences by length and selecting the longest sequence (“length” method). An additional filter can be applied to protein-coding loci, termed “translate”. This is an extension of the “length” method, which limits sequences to those containing a valid reading frame (determined by translation in all forward and reverse frames), thereby removing sequences with errors. However, if no sequences pass translation, the longest sequence is selected rather than excluding the taxon. The “randomize” feature can be used to select a sequence randomly from the set available for a taxon, which will generate supermatrix permutations. Finally, the “vouchered” option will only allow sequences with a voucher tag (generated by *Parse_Loci.py*). For all selection options, sequences must meet a minimum base-pair threshold set by the user. This will determine the smallest amount of data that can be included for a given marker for a given terminal taxon. However, the optimal minimum is another unresolved issue.

To build a population-level dataset, all sequences passing the minimum base pair threshold will be kept. The “translate” option can be used to only include sequences that pass translation, and the “vouchered” option will only include sequences with a voucher tag. The “vouchered” option should be selected to build a population-level dataset that allows samples to be linked by voucher information. Additional information on how various options affect supermatrix and population-level datasets is available online.

The *Filter_Seqs_and_Species.py* module provides key output files for reproducibility and transparency. For each locus, this includes a BatchEntrez-compatible list of all accession numbers from the input file, a per-species list of accession numbers, and a comprehensive summary of the sequence(s) selected for each species (accession number, length, translation test results, and number of alternative sequences available). The *Infer_Supermatrix_Combinations.py* module can be used to infer the total number of possible supermatrix combinations (based on the number of available alternative sequences per taxon per locus). Following the selection of representative sequences, the *Make_Acc_Table.py* module can be used to generate a table of GenBank accession numbers for all taxa and loci. This can be created for species-level supermatrices and “vouchered” population-level datasets.

### 3.7 Multiple Sequence Alignment

SuperCRUNCH includes two pre-alignment steps and several options for multiple sequence alignment (Fig. 4). One pre-alignment module (*Adjust_Direction.py*) adjusts the direction of all sequences in each locus-specific fasta file in combination with mafft. This step produces unaligned fasta files with all sequences written in the correct orientation (thereby avoiding major pitfalls with aligners). Sequences for any locus can be aligned using the *Align.py* module with one of several popular aligners (mafft, muscle, clustal-o) or with all aligners sequentially. For protein-coding loci, the macse translation aligner is also available, which is capable of aligning coding sequences with respect to their translation while allowing for multiple frameshifts or stop codons. To use this alignment method, the *Coding_Translation_Tests.py* module can be used to identify the correct reading frame of sequences, adjust them to the first codon position, and ensure completion of the final codon. Although macse can be run on a single set of reliable sequences (e.g., only those that passed translation), it has an additional feature allowing simultaneous alignment of a set of reliable sequences and a set of unreliable sequences (e.g., those that failed translation), using different parameters. The *Coding_Translation_Tests.py* module can be used to generate all the necessary input files to perform this type of simultaneous alignment using macse (see online documentation).

The alignment methods implemented in SuperCRUNCH are not intended to produce ultra-large alignments containing several thousand sequences. To create ultra-large alignments, we recommend using external alignment methods such as SATé-II (Liu et al., 2012), PASTA (Mirarab et al., 2015), or UPP (Nguyen et al., 2015). We also recommend using UPP to create alignments for loci containing a mix of full-length sequences and short sequence fragments, as these conditions are problematic for many alignment methods (Nguyen et al., 2015).

### 3.8 Post-Alignment Tasks

After multiple sequence alignment, there are several tasks that can be help prepare datasets for downstream analyses. One important task involves relabeling sequences using the *Fasta_Relabel_Seqs.py* module, such that sequence labels are composed of taxon labels, accession numbers, voucher codes, or some combination. The relabeling strategy will depend on the type of dataset being produced (and whether concatenation is intended). Recommendations are provided in the online documentation. Regardless, this step is essential because full-length labels are incompatible with many downstream programs. Relabeled fasta files can be converted into other commonly used formats (nexus, phylip) using the *Fasta_Convert.py* module.

SuperCRUNCH offers two different approaches for automated alignment trimming, although the overall value of trimming remains debatable (Tan et al., 2015). The *Trim_Alignments_Trimal.py* module uses several implementations of trimal (“gap-threshold”, “gappyout”, “noallgaps”) to trim alignments. The *Trim_Alignments_Custom.py* module is based on the custom trimming routine in phyluce (Faircloth, 2016). This version allows edge trimming, row trimming, or both.

Relabeled alignment files can be concatenated using the *Concatenation.py* module. This module allows fasta or phylip input and output formats. The user can also select the symbol for missing data (-, N, ?). It produces a log file containing the number of loci for each terminal taxon and a data partitions file (containing the corresponding base pairs for each locus in the alignment). The *Concatenation.py* module can be used for any dataset in which labels are consistent across loci, including species-level supermatrices (with taxon labels) and “vouchered” population-level datasets (with taxon/voucher combination labels). See online documentation for more details.

## 4 DEMONSTRATIONS AND COMPARISONS

To demonstrate the full range of features available in SuperCRUNCH, we constructed several types of datasets. These included small population-level datasets (<300 sequences, <10 loci), a “vouchered” phylogeographic dataset (∼100 samples, 4 loci), traditional supermatrices (∼1,500 species, ∼70 loci), and phylogenomic supermatrices (∼2,000 UCE loci, <20 samples). In addition, we demonstrate how SuperCRUNCH can be used to add published outgroup sequences to a supermatrix of locally generated sequences. Finally, we compared the ability of SuperCRUNCH to construct species-level supermatrices relative to the program PyPHLAWD (Smith & Walker, 2018), using two test clades (Iguania and Dipsacales). In addition to comparing supermatrix characteristics (taxa, loci, sequences), we also compared the resulting phylogenies (including the number of genera and families recovered as monophyletic). Details are given in Supporting Information S1. All analyses are available as tutorials on the SuperCRUNCH project page on the Open Science Framework (https://osf.io/bpt94/). Analyses were run on an iMac with a 4.2 GHz quad-core Intel Core i7 with 32 GB RAM.

## 5 RESULTS

Detailed results for all analyses are provided in Supporting Information S1, and are briefly summarized here. SuperCRUNCH produced a large supermatrix (∼1,500 species, ∼60 loci, ∼13,000 sequences) in ∼1.5 hours, but with more thorough settings ran up to 13 hours. This difference in runtimes is largely attributable to the alignment step, with MAFFT taking ∼4 minutes and MACSE requiring 11 hours. SuperCRUNCH successfully reconstructed a published phylogeographic dataset (<1 min) and a published phylogenomic supermatrix (∼25 min). It rapidly created new combinations of population-level datasets from multiple published sources (<1 min). It also added GenBank sequences for hundreds of outgroups to a local (unpublished) supermatrix project (<4 min).

SuperCRUNCH outperformed PyPHLAWD in all supermatrix comparisons, recovering more taxa and sequences in both test clades. Given the same starting sequences for the Iguania dataset, SuperCRUNCH found ∼300 more taxa (1,359 vs. 1,069) and ∼2,300 more sequences (12,676 vs. 10,397). PyPHLAWD experienced a severe performance drop for loci containing “complex” records (those with multiple loci or non-overlapping regions), and thereby lost 63% of the available mtDNA sequences (>2,000 sequences discarded). SuperCRUNCH supermatrices also generated higher quality phylogenies, recovering more genera as monophyletic in all comparisons. Additional results for these comparisons are discussed in Supporting Information S1, and all analyses are available on the Open Science Framework (https://osf.io/bpt94/).

## 6 DISCUSSION

SuperCRUNCH is a versatile bioinformatics toolkit that can be used to create large phylogenetic datasets. It contains many novel features that improve its functionality. Most importantly, SuperCRUNCH is not restricted to GenBank sequence data. It can be used to process unpublished sequences, and combinations of GenBank and unpublished data. Many programs rely on GenBank database releases to retrieve starting sequences and obtain metadata. In contrast, SuperCRUNCH infers metadata directly from user-supplied starting sequences, and constructs local databases to perform searches. This design explicitly allows for the inclusion of unpublished sequence data. SuperCRUNCH also includes a key step that allows for either selecting one sequence per species, or all sequences, generating either species-level supermatrices or population-level datasets. Furthermore, filtering options are available for both (passing translation, minimum length), ensuring only high-quality sequences are included in both types of datasets.

A population-level (phylogeographic) dataset includes multiple sequences per species per locus. It is straightforward to collect all sequences available for a particular gene for a given species. However, there may be little overlap of sampling across loci. For example, different individuals may have been sequenced for different loci in different studies. Identifying sequences derived from the same sample can be difficult and requires integrating voucher information. Incorporating additional sequences (published or unpublished) into phylogeographic datasets can be challenging, given the difficulty of identifying and matching voucher information in sequence records. SuperCRUNCH automates these tasks, creating “vouchered” datasets. The “vouchered” feature of SuperCRUNCH only allows sequences with a voucher code to pass the filtering steps used to create a population-level dataset. The final sequences are relabeled using the voucher information (typically taxon name plus voucher code), such that sequences derived from the same sample share an identical label. Together, these features allow the rapid reconstruction of published phylogeographic datasets, merging of published and unpublished data to create new datasets, and construction of datasets from locally generated sequences (especially from sequence-capture experiments).

SuperCRUNCH initially identifies sequences using record labels, moves the relevant sequences to locus-specific files, and performs similarity searches on reduced-sequence sets. Many other programs attempt to cluster all starting sequences to produce putatively orthologous sequence clusters. This can be a useful approach, particularly if the target loci are unknown beforehand. However, these “all-by-all” clustering approaches do not allow target loci to be specified, require additional steps to identify the content of sequence clusters, and can result in the inclusion of paralogous sequences. In certain conditions, the clusters produced from a “complex” record set can be redundant, introducing biases into supermatrices (e.g., a single locus repeated multiple times). SuperCRUNCH putatively assigns sequences to a locus based on the presence of locus search terms in the record label (similar to phyloGenerator). This strategy allows specific loci to be targeted, establishes a clear identity for the sequences, and reduces the chance of including paralogous sequences (which should have a different gene label). In SuperCRUNCH, the label-matching strategy is always paired with downstream similarity searches (e.g. BLASTn). This design provides multiple filters to help eliminate non-target sequences. Thus, SuperCRUNCH can accurately target and build datasets composed of thousands of loci, including UCEs and other sequence-capture loci. It is difficult to reliably perform this task using “all-by-all” clustering of starting sequences. Even the recently proposed “baited” clustering approach of PyPHLAWD, which requires a reference sequence set for each locus, is prohibitive for large genomic datasets (e.g., ∼5,000 UCE loci). However, we acknowledge the success of the label-matching strategy relies on defining appropriate search terms. Unanticipated issues like gene name synonymies can exclude relevant sequences during label-matching (Supporting Information S1). When orthologous sequences are labeled under synonymous gene names, “all-by-all” clustering will recover more sequences if the full set of gene names is not included in the label-matching search (Supporting Information S1). This problem can be partially mitigated by using gene databases to identify synonymies. In SuperCRUNCH, multiple gene abbreviations and gene descriptions can be included to search for a given target locus. Given that searches for loci are conducted using SQL, they are fast and can be executed using iteratively refined search terms to optimize results.

SuperCRUNCH also offers improved methods for similarity searches. These include the ability to specify BLASTn algorithms, improved BLAST coordinate merging and sequence trimming, and flexible choices for selecting reference sequences. Unless specified, the default algorithm used by nucleotide BLAST is megablast, which is best for finding highly similar sequences in intraspecific searches (e.g., population-level datasets). In contrast, discontiguous megablast performs substantially better for interspecific searches (Ma, Tromp, & Li, 2002; Camacho et al., 2009), and is preferable for species-level supermatrices. In many cases, merging the BLAST coordinates obtained from a query sequence is trivial and results in a single continuous target region. However, multiple non-overlapping target regions may also occur for a query sequence, and SuperCRUNCH offers several novel options to handle these cases (Fig. 3). Furthermore, SuperCRUNCH uses the resulting coordinates to automatically trim sequences to the target region, if necessary. This non-standard trimming action ensures that only sequence regions homologous to the reference-sequence set are kept. SuperCRUNCH also offers two options for designating reference sequences: reference sequences can be selected automatically from the sequence set, or can be supplied by the user (Fig. 2). Automatic selection of reference sequences is appropriate for “simple” sequence records (i.e., same gene regions), and can efficiently perform similarity searches for thousands of loci. User-supplied references are more appropriate for “complex” sequence records (multiple loci or non-overlapping regions), or whenever fine-control over the target region is desired. Although this latter option requires gathering reference sequences manually, it is powerful and can be used to extract a single mtDNA gene region from a record set containing a mix of whole mitochondrial genomes, long multi-gene mtDNA sequences, and shorter target sequences.

Despite many improvements implemented in SuperCRUNCH, an important and general issue is the accuracy of GenBank sequence data. This issue can affect SuperCRUNCH and all other programs that process GenBank data. For example, errors may arise through incorrect uploading of data, misidentified specimens, contamination, and other lab errors. Data errors can occur in record labels, and include incorrect gene, taxon, or voucher information. With regards to contamination, we identified two human mtDNA sequences labeled as lizards in our iguanian supermatrix analysis (HM040901.1, KP899454.1; Supplemental File 1). The contamination filter in SuperCRUNCH can detect and eliminate some problems of this kind, but it cannot readily identify cases of misidentified or mislabeled sequences within the focal group. Misidentified specimens are perhaps the most difficult problem to detect, particularly at a shallow taxonomic scale (e.g., a specimen assigned to the wrong species within the same genus or family). Although similarity filtering can generally be used to correctly establish gene identities, parallel approaches for identifying inaccurate taxon labeling within the focal group are generally lacking. Overall, data accuracy is a general problem for the supermatrix approach regardless of the methods used to process the data. Automatic identification of inaccurate sequence records would be a useful goal for future studies of supermatrix construction.

The initial motivation behind SuperCRUNCH was to increase transparency and reproducibility across all steps in dataset construction. We therefore encourage researchers running analyses with SuperCRUNCH to publish the information needed to reproduce their results. This includes accession numbers for the starting sequence set, the taxon list file, the locus search terms file, and the ancillary files and commands used to execute steps. We also emphasize that SuperCRUNCH is highly modular and performing a SuperCRUNCH analysis does not require running the full pipeline. As such, SuperCRUNCH modules can be incorporated into any bioinformatics pipeline or used in conjunction with features of other currently available programs. Alternative programs offer important features that may serve different needs beyond those available in SuperCRUNCH (e.g., SUPERSMART performs phylogenetic analyses on the supermatrices that it generates). Given the rapid growth of sequence data on GenBank (NCBI, 2019) and the changing landscape of phylogenomics, flexible and adaptable bioinformatics approaches are needed to continue mining and managing phylogenetic datasets.

## ACKNOWLEDGMENTS

For financial support, we thank U.S. National Science Foundation Grant DEB 1655690. We thank early testers, including Itzue W. Caviedes-Solis, Cristian Román-Palacios, Benjamin R. Karin, and Pascal O. Title, and participants of the 2019 Trees in the Desert workshop (Tucson, AZ) for their beneficial feedback. We thank Rayna C. Bell, the Bell lab group, the Associate Editor, and three anonymous reviewers for helpful comments that greatly improved the manuscript.

## AUTHORS’ CONTRIBUTIONS

DMP designed the methodology, wrote all code, and analyzed the data; DMP and JJW wrote the manuscript.

## DATA ACCESSIBILITY

SuperCRUNCH is open-source and freely available at https://github.com/dportik/SuperCRUNCH; Zenodo DOI: https://doi.org/10.5281/zenodo.3715362. The complete materials (and instructions for replicating our analyses), including input and output files from each step, is available from the Open Science Framework (https://osf.io/bpt94/).

