## Supporting Information S1 for "SuperCRUNCH: A bioinformatics toolkit for creating and manipulating supermatrices and other large phylogenetic datasets"

**for**

**All datasets produced for this study are available online (with tutorial-style instructions)**

**on the Open Science Framework page for SuperCRUNCH: https://osf.io/bpt94/**

**DEMONSTRATIONS, COMPARISONS, AND RESULTS**

| **Section** | **Page** |
| --- | --- |
| **1 \| *de novo* Supermatrix for Iguania** | **3** |
| 1.1 \| Methods: Iguania-Fast and Iguania-Thorough Analyses | 3 |
| 1.2 \| Results: Iguania-Fast and Iguania-Thorough Analyses | 7 |
| **2 \| UCE Supermatrix for the Genus *Kaloula*** | **11** |
| 2.1 \| Methods: *Kaloula*-Vouchered and *Kaloula*-Species Analyses | 11 |
| 2.2 \| Results: *Kaloula*-Vouchered and *Kaloula*-Species Analyses | 13 |
| **3 \| Phylogeographic Dataset for *Trachylepis sulcata*** | **15** |
| 3.1 \| Methods: *Trachylepis*-Phylogeography and *Trachylepis*-Species Analyses | 15 |
| 3.2 \| Results: *Trachylepis*-Phylogeography and *Trachylepis*-Species Analyses | 16 |
| **4 \| Population Datasets for *Callisaurus* and *Uma*** | **19** |
| 4.1 \| Methods: *Callisaurus*-Population and *Uma*-Population Analyses | 19 |
| 4.2 \| Results: *Callisaurus*-Population and *Uma*-Population Analyses | 20 |
| **5 \| Adding Outgroups to an Unpublished Supermatrix Project: Hyperoliidae** | **23** |
| 5.1 \| Methods: Hyperoliid-Outgroup Analysis | 23 |
| 5.2 \| Results: Hyperoliid-Outgroup Analysis | 25 |
| **6 \| SuperCRUNCH Comparison to PyPHLAWD** | **28** |
| 6.1 \| Methods – PyPHLAWD Comparison | 28 |
| 6.2 \| Results – PyPHLAWD Comparison | 31 |
| 6.2.1\| Supermatrix Results | 31 |
| 6.2.2\| Phylogenetic Results | 34 |
| **7 \| References** | **41** |

**1 | *de novo* Supermatrix for Iguania**

**1.1 | Methods: Iguania-Fast and Iguania-Thorough Analyses**

To demonstrate the full use of SuperCRUNCH we assembled a *de novo* supermatrix for Iguania, a clade of squamate reptiles that contains ~1900 species in 14 families (Uetz et al., 2018). This clade includes chameleons, dragons, flying lizards, iguanas, anoles, and other well-known lizards. For starting material, we downloaded all available iguanian sequence data from GenBank on November 30, 2018 using the organism identifier code for Iguania (txid8511[Organism:exp]). This produced a 13.2 GB fasta file that included 8,785,378 records. We obtained search terms for 69 loci (62 nuclear, 7 mitochondrial) that have been widely used in reptile phylogenetics or phylogeography (Townsend et al., 2008; Portik et al., 2012; Pyron et al., 2013). This includes 62 nuclear loci (*ADNP, AHR, AKAP9, AMEL, BACH1, BACH2, BDNF, BHLHB2, BMP2, CAND1, CARD4, CILP, CMOS, CXCR4, DLL1, DNAH3, ECEL1, ENC1, EXPH5, FSHR, FSTL5, GALR1, GHSR, GPR37, HLCS, INHIBA, KIAA1217, KIAA1549, KIAA2018, KIF24, LRRN1, LZTS1, MC1R, MKL1, MLL3, MSH6, MXRA5, MYH2, NGFB, NKTR, NOS1, NT3, PDC, PNN, PRLR, PTGER4, PTPN, R35, RAG1* (two fragments)*, RAG2, REV3L, RHO, SLC30A1, SLC8A1, SLC8A3, SNCAIP, SOCS5, TRAF6, UBN1, VCPIP, ZEB2, ZFP36L1*) and 7 mitochondrial genes (*12S, 16S, CO1, CYTB, ND1, ND2, ND4*). For the taxon names list, we used a modified version of the February 2018 release of the Reptile Database (which does not contain subspecies). The above starting materials were used to run two different analyses, including one high-quality analysis that used all modules and features (termed “Iguania-Thorough”; https://osf.io/9gs32/), and one analysis that used the fastest possible settings (termed “Iguania-Fast”; https://osf.io/x5hrm/).

For the Iguania-Thorough analysis, we performed an initial taxon assessment of the starting sequences (8,785,378 records). Using *Taxon_Assessment.py*, we found that 158,935 records had a name matching our taxonomy, and 8,626,443 records had a name that did not match a name in our taxonomy. After identifying all correctable unmatched names using the Reptile Database (Uetz et al., 2018), we successfully relabeled 1,860 records using corrected names. We merged the updated records with those that passed the initial taxon assessment, and the resulting sequence set contained 160,795 records. These records were used to search for the 69 loci using *Parse_Loci.py*, and we recovered two or more sequences for all but three loci (*KIAA1217, KIAA1549, MYH2*). We performed similarity filtering using *Cluster_Blast_Extract.py* for 58 nuclear loci (allowing automatic reference selection, intended for “simple” records). We performed similarity filtering using *Reference_Blast_Extract.py* for the 7 mitochondrial genes and for the nuclear protein-coding locus RAG1. This latter filtering module utilizes user-supplied references, and is intended for “complex” record sets (e.g., with little or no overlap for some sequences in some taxa). We included RAG1 in the user-supplied reference strategy because it has been sequenced in (depending on the species) either its entirety as well as for two non-overlapping fragments. We therefore targeted the two regions of RAG1 independently (labeled p1 and p2), which we considered as separate loci downstream. We used two reference sequence sets (*RAG1p1, RAG1p2*) created from the full RAG1 gene of seven tetrapod species (available at https://github.com/dportik/SuperCRUNCH/tree/master/data/reference-sequence-sets/vertebrate-RAG1). To target each of the 7 mitochondrial genes, we assembled a reference sequence set for each gene from 114 squamate mitochondrial genomes, which were downloaded from GenBank. We used the “extract annotation” feature of Geneious (https://www.geneious.com) to quickly obtain each gene region from all mitogenomes (in GenBank format). These reference sequence sets are available at: https://github.com/dportik/SuperCRUNCH/tree/master/data/reference-sequence-sets/squamate-mtdna. After similarity filtering, we selected representative sequences for each species per gene, using options specific to each type of marker (nuclear protein-coding, mtDNA protein-coding, and mtDNA rRNA genes). We used the “oneseq” and “translate” options of *Filter_Seqs_and_Species.py* for the 60 nuclear protein-coding loci (setting translation to standard code) and for the 5 mitochondrial protein-coding genes (setting translation to vertebrate mtDNA code), and the “oneseq” and “length” options for the two mitochondrial rRNA genes (12S, 16S). For each of these three runs, we enforced a minimum 200 bp length for all sequences. We used *Fasta_Filter_by_Min_Seqs.py* to remove alignments with fewer than 30 taxa, which eliminated 6 of the initial 67 loci. We performed sequence direction adjustments for the 61 loci using *Adjust_Direction.py*. We used the *Coding_Translation_Tests.py* module to prepare the 54 nuclear loci (setting translation to standard code) and the 5 mitochondrial protein-coding genes (setting translation to vertebrate mtDNA code) for translation alignment. We used the *Align.py* module to perform MACSE translation alignments, using the “pass_fail” option, for the 54 nuclear loci (setting translation to standard code) and the 5 mitochondrial protein-coding genes (setting translation to vertebrate mtDNA code). We performed sequence alignment for the two rRNA mitochondrial genes (12S, 16S) using Clustal-O using the defaults in *Align.py*. We constructed a table of GenBank accession numbers for all filtered sequences using the *Make_Acc_Table.py* module. We relabeled sequences in all alignment files using the “species” option in *Fasta_Relabel_Seqs.py*, and subsequently trimmed alignments using the gap-threshold option in trimAl with a threshold value of 0.1, as implemented in *Trim_Alignments_Trimal.py*. We converted fasta alignments to phylip and nexus format and concatenated alignments to produce the final supermatrix. In total, we ran 21 separate steps using 16 modules for the Iguania-Thorough analysis (Table S1). The input and output files for all steps, along with complete commands used to execute modules, are available are available on the Open Science Framework at: https://osf.io/9gs32/.

For the Iguania-Fast analysis, we skipped the taxonomy assessment step and began by searching for the 69 loci in the full set of 8,785,378 starting records using *Parse_Loci.py*. We performed similarity filtering using *Cluster_Blast_Extract.py* for 58 nuclear loci (allowing automatic reference selection), and performed similarity filtering using *Reference_Blast_Extract.py* for the 7 mitochondrial genes plus *RAG1* (with the same user-supplied references as above). Following similarity filtering, we selected sequences for all loci using the “length” option in *Filter_Seqs_and_Species.py*, requiring a minimum length of 200 bp. We used *Fasta_Filter_by_Min_Seqs.py* to remove alignments with fewer than 30 taxa, which eliminated 7 of the initial 67 genes. We performed sequence direction adjustments for the remaining 60 loci using *Adjust_Direction.py*, and performed multiple sequence alignment using MAFFT in *Align.py*. We constructed a table of GenBank accession numbers for all filtered sequences using the *Make_Acc_Table.py* module. We relabeled sequences in all alignment files using the “species” option in *Fasta_Relabel_Seqs.py*, and subsequently trimmed alignments using the gap-threshold option in trimAl with a threshold value of 0.1, as implemented in *Trim_Alignments_Trimal.py*. We skipped format conversion and concatenated the alignments to produce the final supermatrix. In total, we ran 11 separate steps using 11 modules for the Iguania-Fast analysis (Table S2). The input and output files for each step, along with complete instructions, are available at: https://osf.io/x5hrm/.

We investigated the quality of the phylogenetic trees resulting from each of the supermatrices (in terms of the number of monophyletic genera, subfamilies, and families). For each supermatrix we ran an unpartitioned RAxML analysis using the GTRCAT model and 100 rapid bootstraps to generate support values. We performed an assessment of these trees in conjunction with other trees of Iguania produced by PyPHLAWD and a constrained SuperCRUNCH analysis (see section 6 – Comparison to PyPHLAWD).

**1.2 | Results: Iguania-Fast and Iguania-Thorough Analyses**

The Iguania-Thorough analysis resulted in a supermatrix containing 61 loci, 1,426 species, and 13,307 total sequences. The analysis took 12 hours and 53 minutes to complete (not including user-time; Table S1). The initial record set contained over 8 million sequences. This set was narrowed down to 160,795 sequences corresponding to the 67 loci. This difference indicates that most sequences downloaded in our GenBank search were irrelevant for our purposes (shotgun genome sequences, mRNA, etc.). Of the 160,795 relevant sequences, 1,860 represent records that were “rescued” by updating an unmatched taxon label. The 160,795 sequences were narrowed down to 13,389 sequences during the sequence-selection step (in which one sequence per taxon per locus was selected). Although the analysis initially found 67 loci, the requirement of at least 30 species per locus eliminated six loci. Consequently, the final number of sequences in the supermatrix totaled 13,307. For the Iguania-Thorough analysis, most steps required only seconds to complete, and a majority of the analysis time is attributable to similarity filtering (~1 hour combined) and multiple sequence alignment (~11.5 hours combined; Table S1). The overall analysis time of ~13 hours was calculated based on running the modules/steps sequentially, but running the three alignment steps simultaneously (MACSE for mtDNA, MACSE for nucDNA, Clustal-O for noncoding mtDNA) would have reduced the total analysis time for Iguania-Thorough to ~7 hours.

The Iguania-Fast analysis resulted in a supermatrix containing 60 loci, 1,399 species, and 12,978 total sequences. The analysis took 1 hour and 28 minutes to complete (not including user-time; Table S2). This analysis did not include any taxonomy assessment, thereby losing the 1,860 records that were “rescued” in the Iguania-Thorough analysis. Rather, loci were parsed directly from the starting set of 8 million records. As a result, the Iguania-Fast analysis recovered less data (1,399 species, 12,978 sequences) than the Iguania-Thorough analysis (1,426 species, 13,307 sequences). This Iguania-Fast analysis also found 67 loci initially, but seven loci were discarded because they contained fewer than 30 species. The Iguania-Fast analysis also required ~1 hour for similarity filtering, but the multiple sequence alignment step using MAFFT took less than 5 minutes to complete.

Most steps of the Iguania-Fast and Iguania-Thorough analyses took similar amounts of time. However, alignment time (5 minutes vs. ~11.5 hours) appears to be the main driver of differences in total time for the Iguania-Fast analysis (~1.5 hours) and the Iguania-Thorough analysis (~13 hours). The Iguania-Thorough analysis resulted in more taxa and sequences, and this was entirely due to the taxonomy assessment step, which only required ~20 minutes to complete (excluding time to identify updated names). We therefore strongly recommend performing the taxonomy assessment step for SuperCRUNCH analyses, as it can result improve dataset quality with minimal computational time.

Trees produced from the Iguania-Thorough and Iguania-Fast datasets are discussed in Section 6 (Comparison to PyPHLAWD).

Table S1. Summary of all steps for the Iguania-Thorough analysis, which took ~13 hours to complete (not including user-time).

| Step | Module | Input Details | Flag Information | Elapsed time |
| --- | --- | --- | --- | --- |
| Assess Taxonomy | Taxa_Assessment.py | 8,785,378 records | --no_subspecies | 0:12:47 |
|  | Rename_Merge.py | Relabeled 1,860 records |  | 0:06:42 |
| Parse Loci | Parse_Loci.py | 69 loci to search, 160,795 records | --no_subspecies | 0:02:41 |
| Similarity Filtering | Cluster_Blast_Extract.py | 58 loci | -b dc-megablast, --max_hits 200, -m span, --threads 4 | 0:18:50 |
|  | Reference_Blast_Extract.py | 9 loci | -b dc-megablast, --max_hits 200, -m span, --threads 4 | 0:48:39 |
|  | Contamination_Filter.py | 7 mtDNA loci | -b megablast | 0:00:06 |
| Sequence Selection | Filter_Seqs_and_Species.py | 60 nuclear coding loci | -s oneseq, -f translate, -m 200, --no_subspecies, --table standard | 0:00:21 |
|  | Filter_Seqs_and_Species.py | 5 mtDNA coding loci | -s oneseq, -f translate, -m 200, --no_subspecies, --table vertmtdna | 0:00:34 |
|  | Filter_Seqs_and_Species.py | 2 mtDNA noncoding loci | -s oneseq, -f length, -m 200, --no_subspecies | 0:00:01 |
|  | Fasta_Filter_by_Min_Seqs.py | 67 loci | --min_seqs 30 | 0:00:01 |
| Sequence Alignment | Adjust_Direction.py | 61 loci | --threads 8 | 0:00:51 |
|  | Coding_Translation_Tests.py | 54 nuclear coding loci | --table standard | 0:00:01 |
|  | Coding_Translation_Tests.py | 5 mtDNA coding loci | --table vertmtdna | 0:00:03 |
|  | Align.py | 2 mtDNA noncoding loci | -a clustalo, --accurate, --threads 4 | 0:27:03 |
|  | Align.py | 5 mtDNA coding loci | -a macse, --table vertmtdna, --mem 10, --pass_fail | 5:30:37 |
|  | Align.py | 54 nuclear coding loci | -a macse, --table standard, --mem 10, --pass_fail | 5:24:27 |
| Post-Alignment | Make_Acc_Table.py | 61 loci |  | 0:00:01 |
|  | Fasta_Relabel_Seqs.py | 61 loci | -r species | 0:00:01 |
|  | Trim_Alignments_Trimal.py | 61 loci | -f fasta, -a gt, --gt 0.1 | 0:00:01 |
|  | Fasta_Convert.py | 61 loci |  | 0:00:01 |
|  | Concatenation.py | 61 loci, 1,426 taxa, 13,307 seqs | --informat fasta, --outformat phylip, -s dash | 0:00:01 |
|  |  |  | **Total elapsed time** | **12:53:49** |

Table S2. Summary of all steps for the Iguania-Fast analysis, which took ~1.5 hours to complete (not including user-time).

| Step | Module | Input details | Flag details | Elapsed time |
| --- | --- | --- | --- | --- |
| Parse Loci | Parse_Loci.py | 69 loci to search; 8,785,378 records | --no_subspecies | 0:17:24 |
| Similarity Filtering | Cluster_Blast_Extract.py | 58 loci | -b dc-megablast, --max_hits 200, -m span, --threads 4 | 0:18:30 |
|  | Reference_Blast_Extract.py | 9 loci | -b dc-megablast, --max_hits 200, -m span, --threads 4 | 0:47:07 |
| Sequence Selection | Filter_Seqs_and_Species.py | 67 loci | -s oneseq, -f length, -m 200, --no_subspecies | 0:00:10 |
|  | Fasta_Filter_by_Min_Seqs.py | 67 loci | --min_seqs 30 | 0:00:01 |
| Sequence Alignment | Adjust_Direction.py | 60 loci | --threads 8 | 0:00:50 |
|  | Align.py | 60 loci | -a mafft, --threads 8 | 0:04:16 |
| Post-Alignment | Make_Acc_Table.py | 60 loci |  | 0:00:01 |
|  | Fasta_Relabel_Seqs.py | 60 loci | -r species | 0:00:01 |
|  | Trim_Alignments_Trimal.py | 60 loci | -f fasta, -a gt, --gt 0.1 | 0:00:01 |
|  | Concatenation.py | 60 loci, 1,399 taxa, 12,978 seqs | --informat fasta, --outformat phylip, -s dash | 0:00:01 |
|  |  |  | **Total elapsed time** | **1:28:22** |

**2 | UCE Supermatrix for the Genus *Kaloula***

**2.1 | Methods: *Kaloula*-Vouchered and *Kaloula*-Species Analyses**

To evaluate the ability of SuperCRUNCH to handle phylogenomic datasets, we attempted to reconstruct the UCE matrix published by Alexander et al. (2017). Their matrix was composed of 14 species in the frog genus *Kaloula* and included a maximum of 1,785 loci per sample. One species (*K. conjuncta*) contained four subspecies, and three taxa were represented by multiple vouchered samples. In total, their dataset included 24 samples, but 6 samples were not identified to species (denoted with “sp.” or “cf.”) and were excluded from our analyses. Therefore, the maximum number of samples we targeted was 18, which represented 14 species. Given the characteristics of this phylogenomic dataset, we performed two separate analyses. For our first analysis, we aimed to construct a “vouchered” UCE supermatrix that would include all 18 samples, which we termed the *Kaloula*-Vouchered analysis (https://osf.io/crzp5/). This analysis was intended to partially reconstruct the full “vouchered” matrix used by the authors for their study. For our second analysis, we aimed to construct a species-level UCE supermatrix that would only include a single representative for each of the 14 species, which we termed the *Kaloula*-Species analysis (https://osf.io/crzp5/). In this analysis, we expected the number of loci to increase for the 3 terminal taxa represented by multiple samples (*K. baleata, K. conjuncta conjuncta*, *K. conjuncta negrosensis*), because sequences would be drawn from multiple samples. We downloaded sequence data from GenBank on January 16, 2019 using the search terms “*Kaloula* ultra conserved element”, which resulted in a 32MB fasta file containing 38,568 records. We generated a locus search terms file from the UCE 5k probe set file (available at https://github.com/faircloth-lab/uce-probe-sets), which targeted 5,041 distinct UCE loci. This general use UCE search terms file is freely available at: https://github.com/dportik/SuperCRUNCH/tree/master/data/locus-search-terms. We created the taxon list directly from the starting sequence set using outputs from *Fasta_Get_Taxa.py*, which resulted in 10 species names and 4 subspecies names.

For the *Kaloula*-Vouchered and the *Kaloula*-Species datasets, we conducted the same general steps for both analyses. Given the taxon list was generated from the starting sequences, we skipped the taxonomy assessment and instead began the analyses by searching for the 5,041 UCE loci in the 38,568 starting records using *Parse_Loci.py*. This produced 1,785 UCE files that each contained more than two sequences. Given that all loci were previously identified and filtered in another pipeline (PHYLUCE; Faircloth, 2016), we did not perform similarity filtering. For the *Kaloula*-Vouchered dataset, we used *Filter_Seqs_and_Species.py* to select sequences with the “allseqs”, “length”, and “vouchered” options, requiring a minimum length of 150 bp. For the *Kaloula*-Species dataset, we used *Filter_Seqs_and_Species.py* to select sequences with the “oneseq” and “length” options, requiring a minimum length of 150 bp. In both datasets, one locus was dropped (for which all sequences were less than 150 bp in length). Accession tables were created using *Make_Acc_Table.py* (with or without the “voucherize” option), sequence directions were adjusted using *Adjust_Direction.py*, and all 1,784 loci were aligned using MAFFT. Sequences were relabeled with *Fasta_Relabel_Seqs.py* using the “species” option (*Kaloula*-Species) or the “species” and “voucherize” options (*Kaloula*-Vouchered), and alignments were concatenated to produce the final supermatrices. In total, we ran 8 separate steps using 8 modules for both the *Kaloula*-Vouchered analysis (Table S3) and the *Kaloula*-Species analysis (Table S4). The input and output files for each step, along with complete instructions, are available on the Open Science Framework for the *Kaloula*-Vouchered analysis (https://osf.io/zxnq8/) and the *Kaloula*-Species analysis (https://osf.io/crzp5/). We investigated whether the supermatrices we constructed resulted in phylogenies concordant with those presented by Alexander et al. (2017). For each supermatrix we ran an unpartitioned RAxML analysis (Stamatakis, 2014), and used the GTRCAT model and 100 rapid bootstraps to generate support values. We compared the resulting topologies to those obtained by Alexander et al. (2017).

**2.2 | Results: *Kaloula*-Vouchered and *Kaloula*-Species Analyses**

The *Kaloula*-Vouchered analysis resulted in a phylogenomic supermatrix containing 1,784 loci, 18 samples, and 28,790 total sequences. The analysis took ~25 minutes to complete (not including user-time; Table S3). The number of UCE loci recovered per individual ranged from 1,276–1,664. The *Kaloula*-Species analysis resulted in a phylogenomic supermatrix containing 1,784 loci, 14 species, and 22,717 total sequences. The analysis took ~20 minutes to complete (not including user-time; Table S4). The number of UCE loci recovered per sample ranged from 1,276–1,777. As expected, the terminal taxa represented by multiple samples displayed an increase in the number of sequences recovered (*K. baleata*: from 1,649 to 1,765; *K. c. conjucta*: from 1,649 to 1,756; *K. c. negrosensis*: from 1,664 to 1,777). The *Kaloula*-Vouchered and *Kaloula*-Species analyses successfully found all 1,785 UCE loci reported by Alexander et al. (2017), but one locus was dropped due to short sequence lengths (all <150 bp). The phylogenies produced from both supermatrices are congruent with results obtained by Alexander et al. (2017). The tree files are available online for the *Kaloula*-Vouchered analysis (https://osf.io/zxnq8/) and the *Kaloula*-Species analysis (https://osf.io/crzp5/)

Table S3. Summary of all steps for the *Kaloula*-Vouchered analysis, which took ~25 minutes to complete (not including user-time).

| Step | Module | Input details | Flag details | Elapsed time |
| --- | --- | --- | --- | --- |
| Parse Loci | Parse_Loci.py | 5,041 loci to search; 38,568 records |  | 0:02:06 |
| Sequence Selection | Filter_Seqs_and_Species.py | 1,785 loci | -s allseqs, -f length, -m 150, --vouchered | 0:01:13 |
|  | Make_Acc_Table.py | 1,784 loci | --voucherize | 0:00:02 |
| Sequence Alignment | Adjust_Direction.py | 1,784 loci | --threads 8 | 0:07:22 |
|  | Align.py | 1,784 loci, mafft | -a mafft, --threads 8 | 0:14:23 |
| Post-Alignment | Fasta_Relabel_Seqs.py | 1,784 loci | -r species, -s, --voucherize | 0:00:03 |
|  | Concatenation.py | 1,784 loci, 18 taxa, 28,790 seqs | --informat fasta, --outformat phylip, -s dash | 0:00:01 |
|  |  |  | **Total elapsed time** | **0:25:10** |

Table S4. Summary of all steps for the *Kaloula*-Species analysis, which took ~20 minutes to complete (not including user-time).

| Step | Module | Input details | Flag details | Elapsed time |
| --- | --- | --- | --- | --- |
| Parse Loci | Parse_Loci.py | 5,041 loci to search; 38,568 records |  | 0:02:05 |
| Sequence Selection | Filter_Seqs_and_Species.py | 1,785 loci | -s oneseq, -f length, -m 150 | 0:00:13 |
|  | Make_Acc_Table.py | 1,784 loci |  | 0:00:02 |
| Sequence Alignment | Adjust_Direction.py | 1,784 loci | --threads 8 | 0:07:03 |
|  | Align.py | 1,784 loci, mafft | -a mafft, --threads 8 | 0:10:25 |
| Post-Alignment | Fasta_Relabel_Seqs.py | 1,784 loci | -r species, -s | 0:00:03 |
|  | Concatenation.py | 1,784 loci, 14 taxa, 22,717 seqs | --informat fasta, --outformat phylip, -s dash | 0:00:01 |
|  |  |  | **Total elapsed time** | **0:19:52** |

**3 | Phylogeographic Dataset for *Trachylepis sulcata***

**3.1 | Methods: *Trachylepis*-Phylogeography and *Trachylepis*-Species Analyses**

To evaluate the ability of SuperCRUNCH to reconstruct phylogeographic datasets from GenBank data, we attempted to reconstruct the dataset of Portik et al. (2011) using their published GenBank sequences. This dataset, which was partially published in Portik et al. (2010), consists of four loci sequenced for 88 samples of the lizard species complex *Trachylepis sulcata*, and several outgroups. We downloaded sequence data from GenBank on July 20, 2019 using the search term “*Trachylepis sulcata*”, which resulted in 442 records (<1 MB in size). We created a locus search terms file specific to the four loci included in Portik et al. (2011): *EXPH5, KIF24, RAG1,* and *ND2*. These are a subset of the loci included in the Iguania supermatrix analyses. We obtained taxon names directly from the starting sequence set using *Fasta_Get_Taxa.py*, and used the outputs to create a taxon list that targeted the focal species (*T. sulcata*) and six outgroup species (*T. aurata, T. punctulata, T. varia, T. variegata, T. vittata,* and *T. wahlbergii*). To reconstruct the phylogeographic dataset of Portik et al. (2011) we ran a “vouchered” analysis (termed *Trachylepis*-Phylogeography), which would include all vouchered samples in the final alignments. For comparison, we also created a species-level supermatrix which would be composed of the seven species (termed *Trachylepis*-Species).

We conducted the same general steps for the *Trachylepis*-Phylogeography and the *Trachylepis*-Species analyses. Given the taxon list was generated from the starting sequences, we skipped the taxonomy assessment and instead began the analyses by searching for the four loci in the 442 starting records using *Parse_Loci.py*. We performed similarity filtering using *Cluster_Blast_Extract.py* for all loci. For the *Trachylepis*-Phylogeography dataset, we used *Filter_Seqs_and_Species.py* to select sequences using the “oneseq”, “length”, and “vouchered” options, requiring a minimum length of 200 bp. For the *Trachylepis*-Species dataset, we used *Filter_Seqs_and_Species.py* to select sequences using the “oneseq” and “length” options, requiring a minimum length of 200 bp. Accession tables were created using *Make_Acc_Table.py* (with or without the “voucherize” option), sequence directions were adjusted using *Adjust_Direction.py*, and all loci were aligned using MAFFT with *Align.py*. Sequences were relabeled with *Fasta_Relabel_Seqs.py* using the “species” option (*Trachylepis*-Species) or the “species” and “voucherize” options (*Trachylepis*-Phylogeography), file formats were converted using *Fasta_Convert.py*, and concatenation was performed using *Concatenation.py*. In total, we ran 10 separate steps using 10 modules for the *Trachylepis*-Phylogeography analysis (Table S5) and the *Trachylepis*-Species analysis (Table S6). The input and output files for each step, along with complete instructions, are available on the Open Science Framework for the *Trachylepis*-Phylogeography analysis (https://osf.io/bgc5z/) and the *Trachylepis*-Species analysis (https://osf.io/umswn/). We investigated if the phylogenies produced from each supermatrix were concordant with results presented by Portik et al. (2011). For each supermatrix we ran an unpartitioned RAxML analysis using the GTRCAT model and 100 rapid bootstraps to generate support values. We compared the resulting topologies to those obtained by Portik et al. (2011).

**3.2 | Results: *Trachylepis*-Phylogeography and *Trachylepis*-Species Analyses**

The *Trachylepis*-Phylogeography analysis successfully reconstructed the phylogeographic dataset of Portik et al. (2011). The analysis found sequences of the four loci (*EXPH5, KIF24, RAG1, ND2*) for all 88 vouchered samples of *Trachylepis sulcata*, which resulted in a total of 326 sequences. In addition, vouchered samples representing several outgroups from Portik et al. (2010) and Portik et al. (2011) were also recovered, including *T. aurata* (2 individuals), *T. punctulata* (3), *T. varia* (6), *T. variegata* (5), *T. vittata* (2), and *T. wahlbergii* (2). This resulted in an additional 74 sequences, and the final concatenated alignment of all four loci (e.g., the phylogeographic supermatrix) contained a total of 108 vouchered samples and 400 sequences. The *Trachylepis*-Phylogeography analysis took 37 seconds to complete (not including user-time; Table S5). The *Trachylepis*-Species analysis was run as a comparison to the *Trachylepis*-Phylogeography analysis. It was used to select one representative sequence per species per locus, for the purpose of creating a species-level matrix from these population-level data. The *Trachylepis*-Species analysis resulted in a matrix containing 7 species (*T. aurata,* *T. punctulata*, *T. sulcata*, *T. varia*, *T. variegata*, *T. vittata*, and *T. wahlbergii*), four loci, and a total of 26 sequences. The *Trachylepis*-Species analysis took 21 seconds to complete (not including user-time; Table S6). The phylogenies produced from the phylogeographic matrix and the species-level matrix were congruent with results presented by Portik et al. (2010) and Portik et al. (2011). The tree files are available online for the *Trachylepis*-Phylogeography analysis (https://osf.io/bgc5z/) and the *Trachylepis*-Species analysis (https://osf.io/umswn/).

Table S5. Summary of all steps for the *Trachylepis*-Phylogeography analysis, which took <1 minute to complete (not including user-time).

| Step | Module | Input details | Flag details | Elapsed time |
| --- | --- | --- | --- | --- |
| Parse Loci | Parse_Loci.py | 4 loci to search; 442 records |  | 0:00:01 |
| Similarity Filtering | Cluster_Blast_Extract.py | 4 loci | -b dc-megablast, -m span, --threads 4 | 0:00:13 |
| Sequence Selection | Filter_Seqs_and_Species.py | 4 loci | -s allseqs, -f length, -m 200, --vouchered | 0:00:01 |
|  | Make_Acc_Table.py | 4 loci | --voucherize | 0:00:01 |
| Sequence Alignment | Adjust_Direction.py | 4 loci | --threads 8 | 0:00:02 |
|  | Align.py | 4 loci | -a mafft, --threads 8 | 0:00:16 |
| Post-Alignment | Fasta_Relabel_Seqs.py | 4 loci | -r species, --voucherize | 0:00:01 |
|  | Fasta_Convert.py | 4 loci |  | 0:00:01 |
|  | Concatenation.py | 4 loci, 108 taxa, 400 seqs | --informat fasta, --outformat phylip, -s dash | 0:00:01 |
|  |  |  | **Total elapsed time** | **0:00:37** |

Table S6. Summary of all steps for the *Trachylepis*-Species analysis, which took <30 seconds to complete (not including user-time).

| Step | Module | Input details | Flag details | Elapsed time |
| --- | --- | --- | --- | --- |
| Parse Loci | Parse_Loci.py | 4 loci to search; 442 records |  | 0:00:01 |
| Similarity Filtering | Cluster_Blast_Extract.py | 4 loci | -b dc-megablast, -m span, --threads 4 | 0:00:13 |
| Sequence Selection | Filter_Seqs_and_Species.py | 4 loci | -s oneseq, -f length, -m 200 | 0:00:01 |
|  | Make_Acc_Table.py | 4 loci |  | 0:00:01 |
| Sequence Alignment | Adjust_Direction.py | 4 loci | --threads 8 | 0:00:01 |
|  | Align.py | 4 loci | -a mafft, --threads 8 | 0:00:01 |
| Post-Alignment | Fasta_Relabel_Seqs.py | 4 loci | -r species | 0:00:01 |
|  | Fasta_Convert.py | 4 loci |  | 0:00:01 |
|  | Concatenation.py | 4 loci, 7 taxa, 26 seqs | --informat fasta, --outformat phylip, -s dash | 0:00:01 |
|  |  |  | **Total elapsed time** | **0:00:21** |

**4 | Population Datasets for *Callisaurus* and *Uma***

**4.1 | Methods: *Callisaurus*-Population and *Uma*-Population Analyses**

We used SuperCRUNCH to generate new combinations of population-level datasets from published sequences. We created independent population-level datasets for the lizard genera *Callisaurus* and *Uma* (family Phrynosomatidae). These were chosen because we knew in advance that some phylogeographic data were present across multiple studies (including Lindell et al., 2005; Schulte & de Queiroz, 2008; Gottscho et al., 2017). However, we did not know which loci would be most strongly represented. We therefore used SuperCRUNCH to survey the availability of sequences. We downloaded sequence data from GenBank on January 25, 2019 using the search term “Phrynosomatidae”, which resulted in a 52MB fasta file containing 82,557 records. We obtained a taxon list directly from the fasta file, which was pruned to contents of each respective genus for separate searches. For *Callisaurus*, this included 11 taxon names (*Callisaurus draconoides, Callisaurus d. bogerti, Callisaurus d. brevipes, Callisaurus d. carmenensis, Callisaurus d. crinitus, Callisaurus d. draconoides, Callisaurus d. inusitanus, Callisaurus d. myurus, Callisaurus d. rhodostictus, Callisaurus d. splendidus, Callisaurus d. ventralis*). For *Uma*, this included 10 taxon names (*Uma exsul, Uma inornata, Uma notata, Uma n. cowlesi, Uma n. notata, Uma n. rufopunctata, Uma paraphygas, Uma rufopunctata, Uma scoparia, Uma s. scoparia*). We recognize that some of the taxon names are synonyms, but we included all names (without corrections) so as to find all available sequences and to obtain counts of sequences for each respective name. We used the same set of 69 locus search terms from our Iguania analysis to perform searches.

We conducted the same general steps for the *Callisaurus*-Population and *Uma*-Population analyses. Given the taxon list was generated from the starting sequences, we skipped the taxonomy assessment and instead began each analysis by searching for the 69 loci in the 82,557 starting records using *Parse_Loci.py*. This resulted in the recovery of 50 loci for *Callisaurus* and 52 loci for *Uma.* We performed similarity filtering using *Cluster_Blast_Extract.py* for all loci. We used *Filter_Seqs_and_Species.py* to select sequences using the “allseqs” and “length” options (but importantly not the “voucher” option), requiring a minimum length of 200 bp. We used *Fasta_Filter_by_Min_Seqs.py* to remove alignments with fewer than 8 sequences, which resulted in the retention of 7 loci for *Callisaurus* and 5 loci for *Uma*. Sequence directions were adjusted using *Adjust_Direction.py*, and all loci were aligned using MAFFT in *Align.py*. Sequences were relabeled with *Fasta_Relabel_Seqs.py* using the “species_acc” option (which is a combination of the taxon name and accession number), and file formats were converted using *Fasta_Convert.py*. In total, we ran 8 separate steps using 8 modules for both the *Callisaurus*-Population analysis (Table S7) and the *Uma*-Population analysis (Table S8). The input and output files for each step, along with complete instructions, are available for the *Callisaurus*-Population analysis at https://osf.io/7gujb/, and for the *Uma*-Population analysis at https://osf.io/e28tu/. For all genes, we ran separate unpartitioned RAxML analyses. We used the GTRCAT model, and performed 100 rapid bootstraps to generate support values for the gene trees.

**4.2 | Results: *Callisaurus*-Population and *Uma*-Population Analyses**

We used SuperCRUNCH to generate new combinations of published sequences, with the intention of finding loci suitable for population-level analyses. We created a population-level dataset for the lizard genus *Callisaurus* and a separate one for the genus *Uma*.

For the *Callisaurus*-Population analysis, we found 7 loci that contained eight or more sequences, including five mitochondrial genes (*12S, n*=8; *CO1*, *n*=8; *CYTB*, *n*=93; *ND1*, *n*=8; *ND2*, *n*=8) and two nuclear loci (*MC1R*, *n*=70; *RAG1*, *n*=70). Among these seven genes, three contained the most sequences (*CYTB, MC1R, RAG1*), each with 70 or more. Across all genes, there were 10 taxa represented (*C. draconoides*, *C. d. bogerti, C. d. brevipes, C. d. carmenensis, C. d. crinitus, C. d. draconoides, C. d. inusitanus, C. d. myurus, C. d. rhodostictus, C. d. splendidud, C. d. ventralis*). The *Callisaurus* analysis took 35 seconds to complete (not including user-time; Table S7).

For the *Uma*-Population analysis, we found 5 genes that contained eight or more sequences, which were all mitochondrial genes (*12S, n*=8; *CO1*, *n*=19; *CYTB*, *n*=191; *ND1*, *n*=8; *ND2, n*=8). Among these genes, *CYTB* contained the greatest number of sequences (*n*=191) by a considerable margin. Across all loci, there were 10 taxa found (*U. exsul, U. inornata, U. notata, U. n. cowlesi, U. n. notata, U. n. rufopunctata, U. paraphygas, U. rufopunctata, U. scoparia, U. s. scoparia*). The *Uma* analysis took 31 seconds to complete (not including user-time; Table S8).

Because sequences in these datasets were not necessarily from vouchered samples, the sequences were labeled using the species name and GenBank accession number. This allowed every sequence within an alignment to have a unique name, but as a result concatenation was not possible. For these datasets, we created a gene tree from each alignment. The individual gene trees produced from the 7 loci for *Callisaurus* and the 5 loci for *Uma* were congruent with phylogenetic results presented by Schulte & de Queiroz (2008), Lindell, Méndez-de la Cruz, and Murphy (2005), and Gottscho et al. (2017). All gene tree files are available online for the *Callisaurus*-Population analysis (https://osf.io/7gujb/) and the *Uma*-Population analysis (https://osf.io/e28tu/).

Table S7. Summary of all steps for the *Callisaurus*-Population analysis, which took under a minute to complete (not including user-time).

| Step | Module | Input details | Flag details | Elapsed time |
| --- | --- | --- | --- | --- |
| Parse Loci | Parse_Loci.py | 69 loci to search; 82,557 records |  | 0:00:05 |
| Similarity Filtering | Cluster_Blast_Extract.py | 50 loci | -b dc-megablast, -m span, --threads 4 | 0:00:09 |
| Sequence Selection | Fasta_Filter_by_Min_Seqs.py | 46 loci | --min_seqs 8 | 0:00:01 |
|  | Filter_Seqs_and_Species.py | 7 loci | -s allseqs, -f length -m 200 | 0:00:01 |
| Sequence Alignment | Adjust_Direction.py | 7 loci | --threads 8 | 0:00:02 |
|  | Align.py | 7 loci, mafft | -a mafft, --threads 8 | 0:00:14 |
| Post-Alignment | Fasta_Relabel_Seqs.py | 7 loci | -r species_acc, -s | 0:00:01 |
|  | Fasta_Convert.py | 7 loci |  | 0:00:01 |
|  |  |  | **Total elapsed time** | **0:00:35** |

Table S8. Summary of all steps for the *Uma*-Population analysis, which took under a minute to complete (not including user-time).

| Step | Module | Input details | Flag details | Elapsed time |
| --- | --- | --- | --- | --- |
| Parse Loci | Parse_Loci.py | 69 loci to search; 82,557 records |  | 0:00:06 |
| Similarity Filtering | Cluster_Blast_Extract.py | 52 loci | -b dc-megablast, -m span, --threads 4 | 0:00:17 |
| Sequence Selection | Fasta_Filter_by_Min_Seqs.py | 45 loci | --min_seqs 8 | 0:00:01 |
|  | Filter_Seqs_and_Species.py | 5 loci | -s allseqs, -f length -m 200 | 0:00:01 |
| Sequence Alignment | Adjust_Direction.py | 5 loci | --threads 8 | 0:00:01 |
|  | Align.py | 5 loci, mafft | -a mafft, --threads 8 | 0:00:03 |
| Post-Alignment | Fasta_Relabel_Seqs.py | 5 loci | -r species_acc, -s | 0:00:01 |
|  | Fasta_Convert.py | 5 loci |  | 0:00:01 |
|  |  |  | **Total elapsed time** | **0:00:31** |

**5 | Adding Outgroups to an Unpublished Supermatrix Project: Family Hyperoliidae**

**5.1 | Methods: Hyperoliid-Outgroup Analysis**

We used SuperCRUNCH to perform a common but sometimes exceedingly difficult task in phylogenetics: adding published outgroup sequences to an unpublished sequencing project. The unpublished sequences were generated as part of Portik et al. (2019), which focused on the systematics of hyperoliid frogs (family Hyperoliidae). These sequences are now available on GenBank (MK497946–MK499204; MK509481–MK509743), but for this demonstration we used the unpublished version of these sequences. This local dataset consisted of six loci sequenced for ~128 species, but many species were represented by multiple vouchered samples. It contains a total of 266 samples. The fasta file for the unpublished dataset contained 1,522 records (1.3MB), and the records were labeled according to the conventions described in the online documentation (https://github.com/dportik/SuperCRUNCH/wiki/2:-Starting-Sequences#ULS). For this analysis, we wanted to add all available GenBank sequence data for the family Arthroleptidae, which is the sister family of Hyperoliidae (the ingroup). We used the search term “Arthroleptidae” to download all available data from GenBank on October 25, 2019, which resulted in a fasta file containing 2,977 records (3MB). For the local sequences, we wanted to treat all samples as distinct (equivalent to a vouchered analysis), whereas for the outgroups we simply wanted to include all possible data for a given species (equivalent to a species-level analysis, in which sequences for a species most likely come from different samples). Within the local sequences, we intentionally labeled the records such that the species names was followed immediately by a museum/field identifier, which allowed us to take advantage of the flexible “subspecies” option in SuperCRUNCH. The subspecies option allows any three-part name to be used, and the third part of a name can contain either a subspecies label or any alphanumerical identifier. We therefore used *Fasta_Get_Taxa.py* with the “numerical” option to obtain a list of “subspecies” from the local sequences that actually contained the museum/field codes instead of a subspecies label (e.g., *Genus species identifier* as opposed to *Genus species subspecies*), and treated these as “subspecies” throughout the analysis. We independently used *Fasta_Get_Taxa.py* to obtain all taxon names in the arthroleptid outgroup sequences, and created a taxon list composed only of species labels (e.g., no subspecies included). The species labels of the arthroleptids (the outgroup) were combined with the “subspecies” labels of the hyperoliids (the ingroup) to create a combined taxon list. We merged the fasta files of the hyperoliid sequences and the arthroleptid sequences to create a single fasta file of sequences. We constructed locus search terms for the 6 loci, which included one mitochondrial gene (*16S*) and five nuclear loci (*FICD, KIAA2013, POMC, RAG1, TYR*). We used these materials to create a custom supermatrix with SuperCRUNCH, which we termed the Hyperoliid-Outgroup analysis.

We wanted to ensure that the sequences obtained for the outgroups closely matched the regions for each gene present in our hyperoliid (local) sequence data. To accomplish this, we used our local sequences as references during similarity filtering. In order to create the six necessary reference files (a locus-specific fasta file composed of only hyperoliid sequences), we ran *Parse_Loci.py* using the hyperoliid fasta file. We then ran *Parse_Loci.py* on the combined hyperoliid and arthroleptid fasta file (containing GenBank and local sequences) to obtain all sequences for each locus, and subsequently ran *Reference_Blast_Extract.py* for each of the six loci using the corresponding hyperoliid reference set. We used *Filter_Seqs_and_Species.py* to select sequences with the “oneseq” and “length” options, requiring a minimum length of 200 bp. Sequence directions were adjusted using *Adjust_Direction.py*, and all loci were aligned with MAFFT using *Align.py*. Sequences were relabeled with *Fasta_Relabel_Seqs.py* using the “species” option with the “subspecies” feature (allowing the hyperoliid sequences to be labeled as *Genus species identifier*). After relabeling, concatenation was performed using *Concatenation.py*. In total, we ran 8 separate steps using 7 modules for the Hyperoliid-Outgroup analysis (Table S9). The input and output files for each step, along with complete instructions, are available at: https://osf.io/q9nyx/. We ran an unpartitioned RAxML analysis on the final supermatrix using the GTRCAT model and 100 rapid bootstraps.

**5.2 | Results: Hyperoliid-Outgroup Analysis**

The Hyperoliid-Outgroup analysis successfully incorporated GenBank sequences (Arthroleptidae) and locally generated sequences (Hyperoliidae) to produce a supermatrix that contained 6 loci, 365 terminals, and 1,724 sequences. The hyperoliid sequences included multiple vouchered samples for many taxa, and the subspecies feature in SuperCRUNCH was used to include museum/field identifiers as the “subspecies” component in their taxon names. This strategy allowed us to successfully include all 266 samples (rather than selecting representative sequences for each of the 128 species). In contrast, for the arthroleptids we simply wanted to obtain the most complete data possible per species (linking vouchers was not relevant). SuperCRUNCH allowed us to target species-level sampling for the outgroup, but population-level sampling for the ingroup. We produced a dataset containing 1,509 sequences and 266 samples for hyperoliids (local), and containing 225 sequences and 99 species for arthroleptids (GenBank). The Hyperoliid-Outgroup analysis took 3 minutes and 45 seconds to complete (not including user-time; Table S9). The phylogeny produced from the Hyperoliid-Outgroup supermatrix is congruent with that estimated by Portik et al. (2019). The tree file is available online (https://osf.io/q9nyx/).

Table S9. Summary of all steps for the Hyperoliid-Outgroup analysis, which took ~4 minutes to complete (not including user-time).

| Step | Module | Input Details | Flag Information | Elapsed time |
| --- | --- | --- | --- | --- |
| Parse Loci | Parse_Loci.py | 6 loci, 4,499 records | (for combined dataset) | 0:00:01 |
|  | Parse_Loci.py | 6 loci, 1,522 records | (for hyperoliid sequences only) | 0:00:01 |
| Similarity Filtering | Reference_Blast_Extract.py | 6 loci | -b dc-megablast, --max_hits 200, -m span, --threads 4 | 0:03:38 |
| Sequence Selection | Filter_Seqs_and_Species.py | 6 loci | -s oneseq, -f length, -m 200 | 0:00:01 |
| Sequence Alignment | Adjust_Direction.py | 6 loci | --threads 8 | 0:00:01 |
|  | Align.py | 6 loci | -a mafft, --threads 8 | 0:00:01 |
| Post-Alignment | Fasta_Relabel_Seqs.py | 6 loci | -r species, -s | 0:00:01 |
|  | Concatenation.py | 6 loci | --informat fasta, --outformat phylip, -s dash | 0:00:01 |
|  |  |  | **Total elapsed time** | **0:03:45** |

**6 | SuperCRUNCH Comparison to PyPHLAWD**

**6.1 | Methods – PyPHLAWD Comparison**

We compared the ability of SuperCRUNCH to construct species-level supermatrices, relative to PyPHLAWD (Smith & Walker, 2018), using two test clades (Iguania and Dipsacales). Importantly, we focused on PyPHLAWD given that Smith & Walker (2018) showed that PyPHLAWD outperformed other methods for supermatrix construction (specifically phyloGenerator, PhyLoTA, and SUPERSMART). Therefore, if our method outperforms PyPHLAWD, then it should represent the state-of-the-art for supermatrix construction. The performance criteria used by Smith & Walker (2018) was the number of taxa and sequences retrieved by each method. We used these criteria as well, but we also considered whether the clades recovered by each method were consistent with current taxonomy. This latter criterion should help detect whether the supermatrices generated by each method tend to yield problematic phylogenetic results. These problematic results will not be apparent simply from the number of taxa and sequences retrieved by each method.

PyPHLAWD retrieves sequences by interfacing directly with a GenBank database release. It was initially designed to produce orthologous sequence clusters using “all-by-all” clustering methods, but it also offers an option to target specific loci by incorporating sets of user-supplied reference sequences (referred to as a “baited” analysis). To search for the same set of 69 loci for Iguania, we performed a “baited” analysis in PyPHLAWD (“v1.0”, Aug 20, 2018 release) with baits consisting of sequences used for the Iguania-Thorough analysis (e.g., mtDNA references) or obtained from them (e.g., the nuclear loci recovered). This PyPHLAWD analysis (termed Iguania-PyPHLAWD) was run using the default settings in the configuration file. All output files for the Iguania-PyPHLAWD analysis are provided at: https://osf.io/vyxj4/. PyPHLAWD does not offer an option to exclude subspecies, and the taxonomy used relies on the NCBI Taxonomy database. As a result, the taxonomy for the PyPHLAWD analysis was incompatible with the taxonomy we obtained from the Reptile Database (Uetz et al., 2018). Therefore, after the analysis we “corrected” the taxonomy by updating names and removing all subspecies in the alignments. As the steps for performing concatenation in PyPHLAWD were unclear, we concatenated the updated alignments using the *Concatenation.py* module in SuperCRUNCH to produce a final supermatrix.

Although we had already run two species-level supermatrix analyses for Iguania using SuperCRUNCH (Iguania-Thorough, Iguania-Fast), these analyses used sequences downloaded from GenBank directly. For these analyses, it would not be possible to determine if differences in the resulting supermatrices (relative to PyPHLAWD) were due to differences in methodology or differences in the starting sequences used. To allow a direct comparison to PyPHLAWD, we used the Iguania sequence set fetched directly by PyPHLAWD (134,028 records) to perform an additional SuperCRUNCH analysis (termed Iguania-Constrained). For this analysis, we performed an initial taxon assessment of the starting sequences and relabeled records using updated names. Locus parsing, similarity filtering, sequence selection, direction adjustment, and alignment used the same options and settings as the Iguania-Fast analysis. We chose the Iguania-Fast settings because several of these steps resembled options in PyPHLAWD, including sequence selection by length, and alignment with MAFFT. We did not remove any loci based on a minimum requirement for the number of sequences (<30) to allow better comparison to PyPHLAWD. Alignments were relabeled, trimmed, and concatenated following the steps outlined in the Iguania-Thorough analysis. In total, we ran 12 separate steps using 12 modules for the Iguania-Constrained analysis (Table S10). The input and output files for each step, along with complete instructions, are available at: https://osf.io/za2ug/.

We further compared the ability of SuperCRUNCH and PyPHLAWD to construct supermatrices by performing additional analyses using the plant clade Dipsacales. This was the example dataset used by Smith and Walker (2018). Following Smith and Walker (2018), we searched for the same four loci (ITS, matK, rbcl, trnL-trnF) using a “baited” analysis in PyPHLAWD (“v1.0”, Aug 20, 2018 release) using their provided bait sets. This PyPHLAWD analysis (termed Dipsacales-PyPHLAWD) was run using the default settings in the configuration file. All output files for the Dipsacales-PyPHLAWD analysis are provided at: https://osf.io/7jqe4/.

We performed a SuperCRUNCH analysis using the Dipsacales sequence set fetched by PyPHLAWD using the GenBank release database (12,348 records), which we termed Dipsacales-Constrained. The taxonomy table produced by PyPHLAWD was used to create a taxon list for this analysis, which included subspecies. The taxonomy table also contained the original description lines for the downloaded sequences, which we examined to create locus search terms for the four loci. We searched for sequences using *Parse_Loci.py*. We performed similarity filtering for each of the four loci using *Reference_Blast_Extract.py,* using the corresponding bait set as the reference sequences. We selected representative sequences per taxon for all loci using the “onseq” and “length” options in *Filter_Seqs_and_Species.py*, requiring a minimum length of 200 bp. We performed sequence direction adjustments for the loci using *Adjust_Direction.py*, and performed multiple sequence alignment using MAFFT in *Align.py*. We constructed a table of GenBank accession numbers for all filtered sequences using the *Make_Acc_Table.py* module. We relabeled sequences in all alignment files using the “species” option and “subspecies” feature in *Fasta_Relabel_Seqs.py*, and subsequently trimmed alignments using the gap-threshold option in trimAl with a threshold value of 0.1, as implemented in *Trim_Alignments_Trimal.py*. We skipped format conversion and concatenated the alignments to produce the final supermatrix. In total, we ran 9 separate steps using 9 modules for the Dipsacales-Constrained analysis (Table S11). The input and output files for each step, along with complete instructions, are available at: https://osf.io/937yu/.

We constructed a total of four supermatrices for Iguania (Iguania-Thorough, Iguania-Fast, Iguania-Constrained, Iguania-PyPHLAWD), and two supermatrices for Dipsacales (Dipsacales-PyPHLAWD, Dipsacales-Comparison). We summarized differences in the content of the supermatrices, including the total number of taxa, loci, and sequences. In addition, we evaluated differences in the resulting phylogenetic trees from these supermatrices. For each supermatrix we ran an unpartitioned RAxML analysis, and used the GTRCAT model and 100 rapid bootstraps to generate a phylogeny with support values. We evaluated the number of genera, subfamilies, and families recovered as monophyletic in the four different Iguania trees, and the number of monophyletic genera in the two different Dipsacales trees. All tree files are available at: https://osf.io/vgwu3/.

**6.2 | Results – PyPHLAWD Comparison**

**6.2.1 | Supermatrix Results**

SuperCRUNCH generally outperformed PyPHLAWD in all comparisons and resulted in higher numbers of sequences and taxa for the Iguania and Dipsacales datasets (Tables S12–S14). The analysis of Dipsacales was smaller in scope. SuperCRUNCH obtained better results, but the supermatrices generated by each method were generally similar (versus iguanians, see below). For Dipsacales, we successfully replicated the results of Smith and Walker (2018) using PyPHLAWD in terms of the number of species recovered. The Dipsacales-PyPHLAWD analysis resulted in a supermatrix containing 4 loci, 641 taxa, and 1,510 total sequences (Table S14), and took 1 minute and 16 seconds to complete. The Dipsacales-Constrained analysis in SuperCRUNCH resulted in a supermatrix containing 4 loci, 651 taxa, and 1,589 total sequences, and took 1 minute and 14 seconds to complete (not including user-time; Table S11). The Dipsacales-Constrained analysis recovered more sequences for 3 out of the 4 loci, and both analyses recovered an equal number of sequences for 1 locus (Table S14).

The Iguania-PyPHLAWD analysis resulted in a supermatrix containing 66 loci, 1,069 species, and 10,397 total sequences (Table S13), and took 18 minutes to complete. The Iguania-Constrained analysis in SuperCRUNCH generated a supermatrix containing 67 loci, 1,359 taxa, and 12,676 total sequences (Table S13). The analysis took 1 hour and 8 minutes to complete (not including user-time; Table S10). Among the 66 loci shared between the two analyses, the Iguania-PyPHLAWD analysis recovered more sequences for 5 loci (7%), the Iguania-Constrained analysis recovered more sequences for 19 loci (29%), and both analyses recovered equal numbers of sequences for 43 loci (64%; Table S12). These two analyses relied on starting sequences obtained through a GenBank database release. They were outperformed by all other SuperCRUNCH analyses of Iguania that used data downloaded from GenBank directly by us (including Iguania-Thorough and Iguania-Fast; Table S13). In particular, the Iguania-Thorough analysis in SuperCRUNCH vastly outperformed the Iguania-PyPHLAWD analysis. It recovered more species (1,426 vs. 1,069) and total sequences (13,307 vs. 10,397) despite having fewer loci in the final supermatrix (it initially found 67 loci but discarded 6 because of the minimum sequence filter).

A per-locus comparison for Iguania revealed the largest difference in performance between the PyPHLAWD analysis and any of the three SuperCRUNCH analyses was for loci containing “complex” records (those consisting of multiple loci or non-overlapping regions). These included the mitochondrial genes, as well as *RAG1* (Table S12). Given the same set of starting sequences, PyPHLAWD found only 37% of the total mtDNA sequences recovered by SuperCRUNCH in the Iguania-Constrained analysis, and recovered only four of the seven mitochondrial genes with reasonable success (>50 sequences).

The likely source of this issue in PyPHLAWD is the close match required between baits and sequences, which is related to both length and sequence divergence (S. Smith, personal communication). PyPHLAWD (“v1.0”) does not automatically trim query sequences after similarity searches. Rather, it passes or fails an entire sequence based on set threshold values (such as percent identity and minimum or maximum length). In the case of “complex” mtDNA records, the entire multigene sequence would pass or fail, rather than being trimmed to the target region. It may have been possible for us to obtain better results by changing these default settings. Given this design and the sequence length heterogeneity present in “complex” records, it is unclear if there is an optimal setting that would allow all target sequences to be found for these types of loci. Multiple runs using different settings would likely be necessary. Inspection of the similarity searching outputs from SuperCRUNCH confirms this idea (i.e., outputs containing the starting lengths, BLAST coordinates, and trimmed lengths of input sequences). In the case of *CO1* (for which no sequences were recovered with PyPHLAWD), we used several reference sequences spanning the entire length of the *CO1* gene (~1,500 bp) as the “baits”. All input sequences containing *CO1* fell into two categories: (1) they were purely on target but <700 bp in length, or (2) they were “complex” records in which *CO1* represented less than 50% of the length of the total sequence. Both of these scenarios appeared to cause severe problems for the similarity searches in PyPHLAWD (particularly the matched length requirement between bait and sequence), and the default settings caused both categories of sequences to fail this filter. Although we might have obtained better results for one category of sequences for *CO1* by using different settings, this would have driven losses for the other category of sequences. However, even if all sequences were retained, the recovery of multigene *CO1* sequences would result in redundancy in the final supermatrix (multiple mtDNA gene regions present for *CO1*). Given that the characteristics of the input sequences are generally unknown (e.g., “simple” vs. “complex”, length heterogeneity), this makes finding appropriate “baits” and defining appropriate settings challenging. Thus, significant losses of data for some loci are expected to occur with PyPHLAWD (particularly those containing “complex” records), as we observe here.

The results for the nuclear loci (e.g., “simple” records) were much more similar between methods, and both methods obtained the same number of sequences for 43 loci. In some cases, PyPHLAWD obtained more sequences for nuclear loci. Two of these cases revealed a limitation of the label-searching method of SuperCRUNCH, as synonomous gene names for the two loci (R35 vs. GPR14, ZEB2 vs. ZHFX1b) resulted in the loss of actual homologous sequences by SuperCRUNCH. However, we also identified at least one case in which PyPHLAWD included paralogous sequence data (ENC1 contained ENC6 sequences), which artificially inflated the total number of sequences for that locus. We did not observe these paralogous sequences (of ENC6) in the ENC1 alignment created using SuperCRUNCH. Instead, these paralogous sequences were eliminated because they did not contain the gene abbreviation or description terms for ENC1. The results for the nuclear loci therefore revealed important trade-offs associated with the “all-by-all” clustering approach of PyPHLAWD, and the label-searching plus similarity filtering strategy of SuperCRUNCH.

**6.2.2 | Phylogenetic Results**

We compared phylogenetic trees estimated from the supermatrices for the four analyses of Iguania, and the two analyses of Dipsacales. For each iguanian phylogeny, we evaluated the number of genera, subfamilies, and families recovered as monophyletic. The Iguania-Thorough, Iguania-Fast, and Iguania-Constrained analyses recovered all 14 families as monophyletic with high support, whereas Iguania-PyPHLAWD only recovered 13 of 14 families as monophyletic. The PyPHLAWD analysis failed to support the monophyly of Agamidae (i.e., Chamaeleonidae is nested inside Agamidae). Agamid monophyly has been strongly supported in previous supermatrix analyses (Pyron et al. 2013; Zheng & Wiens, 2016) and phylogenomic analyses (Streicher, Schulte, & Wiens 2016). All four Iguania analyses recovered all 10 subfamilies as monophyletic with high support (all >95%). Comparisons of genera were more complex, as the trees each contained a different number of species and genera. The Iguania-Thorough phylogeny contained 111 genera, including 68 monophyletic genera, 16 non-monophyletic genera, and 27 genera represented by a single species. The Iguania-Fast phylogeny contained 108 genera, including 65 monophyletic genera, 15 non-monophyletic genera, and 28 genera represented by a single species. The Iguania-Constrained phylogeny contained 106 genera, including 64 monophyletic genera, 13 non-monophyletic genera, and 29 genera represented by a single species. The Iguania-PyPHLAWD phylogeny contained 94 genera, including 58 monophyletic genera, 12 non-monophyletic genera, and 24 genera represented by a single species. Although PyPHLAWD and SuperCRUNCH produced somewhat similar scores for these metrics, the Iguania-Thorough phylogeny contained an additional 357 taxa and 17 genera not present in the PyPHLAWD phylogeny. The phylogenies and corresponding assessments of the groupings present are provided at: https://osf.io/zxhby/.

For the plant family Dipsacales, we compared the monophyly of genera. In this regard, the Dipascales phylogeny produced from the SuperCRUNCH supermatrix was also higher quality than the phylogeny from the PyPHLAWD supermatrix. These trees contained different numbers of species, but the same number of genera. The Dipsacales-Constrained phylogeny from SuperCRUNCH contained 42 genera, including 24 monophyletic genera, 9 non-monophyletic genera, and 9 genera represented by a single species. The Dipsacales-PyPHLAWD phylogeny contained 42 genera, including 20 monophyletic genera, 11 non-monophyletic genera, and 11 genera represented by a single species. The phylogenies and corresponding assessments of the groupings present are provided at: https://osf.io/zxhby/.

Table S10. Summary of all steps for the Iguania-Constrained analysis, which took ~1 hour to complete (not including user-time).

| Step | Module | Input Details | Flag Information | Elapsed time |
| --- | --- | --- | --- | --- |
| Assess Taxonomy | Taxa_Assessment.py | 134,028 records | --no_subspecies | 0:00:10 |
|  | Rename_Merge.py | Relabeled 430 records |  | 0:00:01 |
| Parse Loci | Parse_Loci.py | 69 loci to search, 133,962 records | --no_subspecies | 0:00:25 |
| Similarity Filtering | Cluster_Blast_Extract.py | 58 loci | -b dc-megablast, --max_hits 200, -m span, --threads 4 | 0:18:04 |
|  | Reference_Blast_Extract.py | 9 loci | -b dc-megablast, --max_hits 200, -m span, --threads 4 | 0:44:41 |
| Sequence Selection | Filter_Seqs_and_Species.py | 67 loci | -s oneseq, -f length, -m 200, --no_subspecies | 0:00:10 |
| Sequence Alignment | Adjust_Direction.py | 67 loci | --threads 8 | 0:00:56 |
|  | Align.py | 67 loci | -a mafft, --threads 4 | 0:04:27 |
| Post-Alignment | Make_Acc_Table.py | 67 loci |  | 0:00:01 |
|  | Fasta_Relabel_Seqs.py | 67 loci | -r species | 0:00:01 |
|  | Trim_Alignments_Trimal.py | 67 loci | -f fasta, -a gt | 0:00:01 |
|  | Concatenation.py | 67 loci, 1,359 taxa, 12,676 seqs | --informat fasta, --outformat phylip, -s dash | 0:00:01 |
|  |  |  | **Total elapsed time** | **1:08:58** |

Table S11. Summary of all steps for the Dipsacales-Constrained analysis, which took ~1 minute to complete (not including user-time).

| Step | Module | Input Details | Flag Information | Elapsed time |
| --- | --- | --- | --- | --- |
| Parse Loci | Parse_Loci.py | 4 loci to search, 12,348 records |  | 0:00:01 |
| Similarity Filtering | Reference_Blast_Extract.py | 4 loci | -b dc-megablast, -m span, --threads 4 | 0:00:26 |
| Sequence Selection | Filter_Seqs_and_Species.py | 4 loci | -s oneseq, -f length, -m 200 | 0:00:01 |
| Sequence Alignment | Adjust_Direction.py | 4 loci | --threads 8 | 0:00:07 |
|  | Align.py | 4 loci | -a mafft, --accurate, --threads 8 | 0:00:35 |
| Post-Alignment | Make_Acc_Table.py | 4 loci | -s | 0:00:01 |
|  | Fasta_Relabel_Seqs.py | 4 loci | -r species, -s | 0:00:01 |
|  | Trim_Alignments_Trimal.py | 4 loci | -f fasta, -a gt, --gt 0.1 | 0:00:01 |
|  | Concatenation.py | 4 loci, 651 taxa, 1,589 seqs | --informat fasta, --outformat phylip, -s dash | 0:00:01 |
|  |  |  | **Total elapsed time** | **0:01:14** |

Table S12. The total number of sequences for each locus in the supermatrix produced by each analysis for the clade Iguania.

| Locus Type | Locus Name | Iguania-PyPHLAWD | Iguania-Constrained | Iguania-Fast | Iguania-Thorough |
| --- | --- | --- | --- | --- | --- |
| mtDNA | 12S | 325 | 489 | 499 | 530 |
|  | 16S | 9 | 544 | 560 | 580 |
|  | CO1 | - | 353 | 482 | 481 |
|  | CYTB | 348 | 494 | 508 | 516 |
|  | ND1 | 109 | 203 | 208 | 209 |
|  | ND2 | 523 | 928 | 945 | 1006 |
|  | ND4 | 1 | 504 | 540 | 552 |
| Nuclear | ADNP | 142 | 142 | 142 | 145 |
|  | AHR | 133 | 133 | 133 | 137 |
|  | AKAP9 | 177 | 180 | 180 | 184 |
|  | AMEL | 10 | 13 | <30 | <30 |
|  | BACH1 | 183 | 183 | 183 | 187 |
|  | BACH2 | 29 | 30 | 30 | 31 |
|  | BDNF | 375 | 379 | 380 | 394 |
|  | BHLHB2 | 146 | 146 | 146 | 149 |
|  | BMP2 | 140 | 140 | 140 | 143 |
|  | CAND1 | 145 | 145 | 145 | 149 |
|  | CARD4 | 139 | 139 | 139 | 141 |
|  | CILP | 143 | 143 | 143 | 145 |
|  | CMOS | 443 | 439 | 447 | 463 |
|  | CXCR4 | 143 | 143 | 143 | 145 |
|  | DLL1 | 140 | 140 | 140 | 140 |
|  | DNAH3 | 137 | 137 | 137 | 141 |
|  | ECEL1 | 150 | 150 | 150 | 150 |
|  | ENC1 | 94 | 41 | 41 | 45 |
|  | EXPH5 | 148 | 148 | 148 | 149 |
|  | FSHR | 145 | 145 | 145 | 148 |
|  | FSTL5 | 141 | 141 | 141 | 144 |
|  | GALR1 | 137 | 137 | 137 | 138 |
|  | GHSR | 132 | 132 | 132 | 132 |
|  | GPR37 | 140 | 140 | 140 | 144 |
|  | HLCS | 137 | 137 | 137 | 139 |
|  | INHIBA | 140 | 140 | 141 | 145 |
|  | KIAA1217 | - | - | - | - |
|  | KIAA1549 | - | - | - | - |
|  | KIAA2018 | 15 | 15 | <30 | <30 |
|  | KIF24 | 139 | 139 | 139 | 139 |
|  | LRRN1 | 132 | 132 | 132 | 132 |
|  | LZTS1 | 122 | 122 | 122 | 122 |
|  | MC1R | 40 | 40 | 40 | 40 |
|  | MKL1 | 114 | 114 | 151 | 153 |
|  | MLL3 | 123 | 123 | 123 | 123 |
|  | MSH6 | 163 | 164 | 164 | 168 |
|  | MXRA5 | 104 | 104 | 107 | 107 |
|  | MYH2 | - | - | - | - |
|  | NGFB | 169 | 169 | 169 | 173 |
|  | NKTR | 215 | 221 | 221 | 227 |
|  | NOS1 | 9 | 9 | <30 | <30 |
|  | NT3 | 350 | 355 | 365 | 368 |
|  | PDC | 19 | 19 | <30 | <30 |
|  | PNN | 269 | 269 | 274 | 277 |
|  | PRLR | 420 | 420 | 438 | 440 |
|  | PTGER4 | 147 | 147 | 147 | 149 |
|  | PTPN | 114 | 114 | 115 | 115 |
|  | R35 | 300 | 288 | 299 | 303 |
|  | RAG1p1 | 344 | 429 | 439 | 455 |
|  | RAG1p2 | 290 | 361 | 390 | 397 |
|  | RAG2 | - | 2 | <30 | <30 |
|  | REV3L | 28 | 29 | <30 | 30 |
|  | RHO | 38 | 13 | 58 | 58 |
|  | SLC30A1 | 144 | 144 | 144 | 147 |
|  | SLC8A1 | 144 | 144 | 144 | 148 |
|  | SLC8A3 | 143 | 143 | 143 | 146 |
|  | SNCAIP | 249 | 249 | 249 | 252 |
|  | SOCS5 | 17 | 17 | <30 | 152 |
|  | TRAF6 | 149 | 149 | 149 | 151 |
|  | UBN1 | 147 | 148 | 148 | 151 |
|  | VCPIP | 131 | 131 | 131 | 133 |
|  | ZEB2 | 164 | 154 | 154 | 157 |
|  | ZFP36L1 | 141 | 141 | 141 | 143 |

Table S13. The total number of loci, taxa, sequences, and the concatenated alignment length produced by each analysis for the clade Iguania.

| Analysis | Loci | Species | Genera | Sequences | Alignment length |
| --- | --- | --- | --- | --- | --- |
| Iguania-PyPHLAWD | 66 | 1,069 | 94 | 10,397 | 66,100 bp |
| Iguania-Constrained | 67 | 1,359 | 106 | 12,676 | 58,315 bp |
| Iguania-Fast | 60* | 1,399 | 108 | 12,978 | 52,827 bp |
| Iguania-Thorough | 61* | 1,426 | 111 | 13,307 | 53,319 bp |

*Note that an additional filter that required a minimum of 30 taxa per locus removed several loci for Iguania-Fast (*n*=7) and Iguania-Thorough (*n*=6).

Table S14. The total number of sequences for each locus in the supermatrix produced by each analysis for the clade Dipsacales.

| Locus | Dipsacales-PyPHLAWD | Dipsacales-Constrained |
| --- | --- | --- |
| ITS | 556 | 583 |
| matK | 322 | 344 |
| rbcl | 299 | 299 |
| trnL-trnF | 333 | 363 |

**7 | References**

Alexander, A. A., Su, Y.-C., Oliveros, C. H., Olson, K. V., Travers, S. L., & Brown, R. M. (2016). Genomic data reveals potential for hybridization, introgression, and incomplete lineage sorting to confound phylogenetic relationships in an adaptive radiation of narrow-mouth frogs. *Evolution, 71*, 475–488.

Faircloth, B. C. (2016). PHYLUCE is a software package for the analysis of conserved genomic loci. *Bioinformatics*, *32*, 786–788.

Gottscho, A. D., Wood, D. A., Vandergast, A. G., Lemos-Espinal, J., Gatesy, J., & Reeder, T. W. (2017). Lineage diversification of fringe-toed lizards (Phrynosomatidae: *Uma notata* complex) in the Colorado Desert: delimiting species in the presence of gene flow. *Molecular Phylogenetics and Evolution*, *106*, 103–117.

Lindell, J., Méndez-de la Cruz, F. R., & Murphy, R. W. (2005). Deep genealogical history without population differentiation: discordance between mtDNA and allozyme divergence in the zebra-tailed lizard (*Callisaurus draconoides*). *Molecular Phylogenetics and Evolution, 36,* 682–694.

Portik, D. M., Bauer, A. M., & Jackman, T. R. (2010). The phylogenetic affinities of *Trachylepis sulcata nigra* and the intraspecific evolution of coastal melanism in the western rock skink. *African Zoology*, 45, 147–159.

Portik, D. M., Bauer, A. M., & Jackman, T. R. (2011). Bridging the gap: Western rock skinks (*Trachylepis sulcata*) have a short history in South Africa. *Molecular Ecology,* *20*, 1744−1758.

Portik, D. M., Wood Jr., P. L., Grismer, J. L., Stanley, E. L., & Jackman, T. R. (2012). Identification of 104 rapidly-evolving nuclear protein-coding markers for amplification across scaled reptiles using genomic resources. *Conservation Genetics Resources, 4,* 1−10.

Portik, D. M., Bell, R. C., Blackburn, D. C., Bauer, A. M., Barratt, C. D., Branch, W. R., Burger, M., Channing, A., Colston, T. J., Conradie, W., Dehling, J. M., Drewes, R. C., Ernst, R., Greenbaum, E., Gvoždík, V., Harvey, J., Hillers, A., Hirschfeld, M., Jongsma, G. F. M., Kielgast, J., Kouete, M. T., Lawson, L., Leaché, A. D., Loader, S. P., Lötters, S., van der Meijden, A., Menegon, M., Müller, S., Nagy, Z. T., Ofori-Boateng, C., Ohler, A., Papenfuss, T. J., Rößler, D., Sinsch, U., Rödel, M. -O., Veith, M., Vindum, J., Zassi-Boulou, A. -G., & McGuire, J. A. (2019). Sexual dichromatism drives diversification within a major radiation of African amphibians. *Systematic Biology*, 68, 859–875.

Pyron, R. A., Burbrink, F. T., & Wiens, J. J. (2013). A phylogeny and revised classification of Squamata, including 4161 species of lizards and snakes. *BMC Evolutionary Biology, 13,* 93.

Schulte II, J. A., & de Queiroz, K. (2008). Phylogenetic relationships and heterogeneous evolutionary processes among phrynosomatine sand lizards (Squamata, Iguanidae) revisited. *Molecular Phylogenetics and Evolution, 47,* 700–716.

Smith, S. A., & Walker, J. F. (2018). PyPHLAWD: A python tool for phylogenetic dataset construction. *Methods in Ecology and Evolution*, 10, 104–108.

Stamatakis, A. (2014). RAxML version 8: a tool for phylogenetic analysis and post-analysis of large phylogenies. *Bioinformatics, 30,* 1312–1313.

Streicher, J. W., Schulte, J. A., & Wiens, J. J. (2016). How should genes and taxa be sampled for phylogenomic analyses with missing data? An empirical study in iguanian lizards. *Systematic Biology,* *65,* 128–145.

Townsend, T. M., Alegre, R. E., Kelley, S. T., Wiens, J. J., & Reeder, T. W. (2008). Rapid development of multiple nuclear loci for phylogenetic analysis using genomic resources: an example from squamate reptiles. *Molecular Phylogenetics and Evolution, 47,* 129–142.

Uetz, P., Freed, P., & Hošek, J. (2018). The Reptile Database. Available at: http://www.reptile-database.org.

Zheng, Y., & Wiens, J. J. (2016). Combining phylogenomic and supermatrix approaches, and a time-calibrated phylogeny for squamate reptiles (lizards and snakes) based on 52 genes and 4,162 species. *Molecular Phylogenetics and Evolution,* *94,* 537–547.
